# A key linear epitope for a potent neutralizing antibody to SARS-CoV-2 S-RBD

**DOI:** 10.1101/2020.09.11.292631

**Authors:** Tingting Li, Xiaojian Han, Yingming Wang, Chenjian Gu, Jianwei Wang, Chao Hu, Shenglong Li, Kai Wang, Feiyang Luo, Jingjing Huang, Yingyi Long, Shuyi Song, Wang Wang, Jie Hu, Ruixin Wu, Song Mu, Yanan Hao, Qian Chen, Fengxia Gao, Meiying Shen, Shunhua Long, Fang Gong, Luo Li, Yang Wu, Wei Xu, Xia Cai, Di Qu, Zhenghong Yuan, Qingzhu Gao, Guiji Zhang, Changlong He, Yaru Nai, Kun Deng, Li Du, Ni Tang, Youhua Xie, Ailong Huang, Aishun Jin

**Author notes:** These authors contributed equally to this work (Tingting Li, Xiaojian Han, Yingming Wang, Chenjian Gu). Correspondence (A.S.J); (Y.H.X); (A.L.H).

## Abstract

The spread of SARS-CoV-2 confers a serious threat to the public health without effective intervention strategies^1–3^. Its variant carrying mutated Spike (S) protein D614G (S^D614G^) has become the most prevalent form in the current global pandemic^4,5^. We have identified a large panel of potential neutralizing antibodies (NAbs) targeting the receptor-binding domain (RBD) of SARS-CoV-2 S^6^. Here, we focused on the top 20 potential NAbs for the mechanism study. Of them, the top 4 NAbs could individually neutralize both authentic SARS-CoV-2 and S^D614G^ pseudovirus efficiently. Our epitope mapping revealed that 16/20 potent NAbs overlapped the same steric epitope. Excitingly, we found that one of these potent NAbs (58G6) exclusively bound to a linear epitope on S-RBD (termed as 58G6e), and the interaction of 58G6e and the recombinant ACE2 could be blocked by 58G6. We confirmed that 58G6e represented a key site of vulnerability on S-RBD and it could positively react with COVID-19 convalescent patients’ plasma. We are the first, as far as we know, to provide direct evidences of a linear epitope that can be recognized by a potent NAb against SARS-CoV-2 S-RBD. This study paves the way for the applications of these NAbs and the potential safe and effective vaccine design.

## Introduction

Pandemic outbreak of COVID-19 has caused high mortality and unprecedented social and economic consequences^1–3^. Recently, researchers have found that a SARS-CoV-2 mutation (S^D614G^) renders easier virus entry and causes advanced infectivity and excess mortality^4,5,7,8^. As SARS-CoV, SARS-CoV-2 utilizes the S-RBD to bind to the same host receptor angiotensin-converting enzyme 2 (ACE2)^9–11^. The structure of SARS-CoV-2 S-RBD-ACE2 complex has been elucidated, which provides fundamental information for the development of NAbs and the design of vaccines^12–14^. Due to the fact that clinically available NAbs and vaccines are currently under investigations, their safety and efficacy remain largely unknown. Independently, our group and others have reported that NAbs could efficiently block authentic virus entry by interfering with SARS-CoV-2 S-RBD and its receptor binding^6,15–19^. Several studies have demonstrated the epitope mapping of the NAbs to SARS-CoV-2 in three dimensions, which has been proved to promote the development of effective vaccines for COVID-19^15–18^. In this study, we exhibited several neutralizing monoclonal antibodies (mAbs) with high potency against authentic SARS-CoV-2 and SARS-CoV-2 S^D614G^ pseudovirus. More importantly, we discovered a linear epitope that could be recognized by a potent NAb, possibly representing a vulnerability site on SARS-CoV-2 S-RBD. The identification of this novel mechanism of NAbs might provide fundamental information for the NAb-based therapies and vaccine design.

## Results

### Characteristics of the binding of potential NAbs to SARS-CoV-2 S or S^D614G^

Previously, we screened a total of 219 mAbs to SARS-CoV-2 S-RBD from the memory B cells of COVID-19 convalescent patients via our established NAbs screening system, and discovered a panel of potential NAbs by the authentic virus cytopathic effect (CPE) assay^6^. The top 20 of these potential NAbs were used for functional characterization. At first, we investigated the binding ability of these Abs to SARS-CoV-2 S protein subunit S1, S-RBD and S^D614G^ The half-maximum efficient concentration (EC_50_) value of these potential NAbs ranged from 13.36 ng/mL to 94.62 ng/mL for the binding to SARS-CoV-2 S1, 23 ng/mL to 131.4 ng/mL to S-RBD, and 17.9 ng/mL to 191.4 ng/mL to S^D614G^ (Fig. 1a, Extended Data Fig. 1, Table 1). We also found that these potential NAbs bound to SARS-CoV only weakly, confirming their specificity to SARS-CoV-2 (data not shown). These results showed that although the binding ability of these potential NAbs varied individually, their capabilities of interacting with SARS-CoV-2 S1, S-RBD and S^D614G^ were similar. Next, we detected the affinity of these 20 Abs by the surface plasmon resonance (SPR) assay. We found that the affinity constant (K_D_) between the RBD and these Abs varied from 0.08 to 8 nM, except for 81C3, with the affinity of 24 nM (Fig. 1a, Extended Data Fig. 2, Table 1). Therefore, we have demonstrated that our top 20 potential NAbs could strongly bind to not only SARS-CoV-2 S and S-RBD, but also S^D614G^, as the most prevalent form of the current SARS-CoV-2 variants, confirming their exploitable value as NAb candidates.

**Fig. 1.**
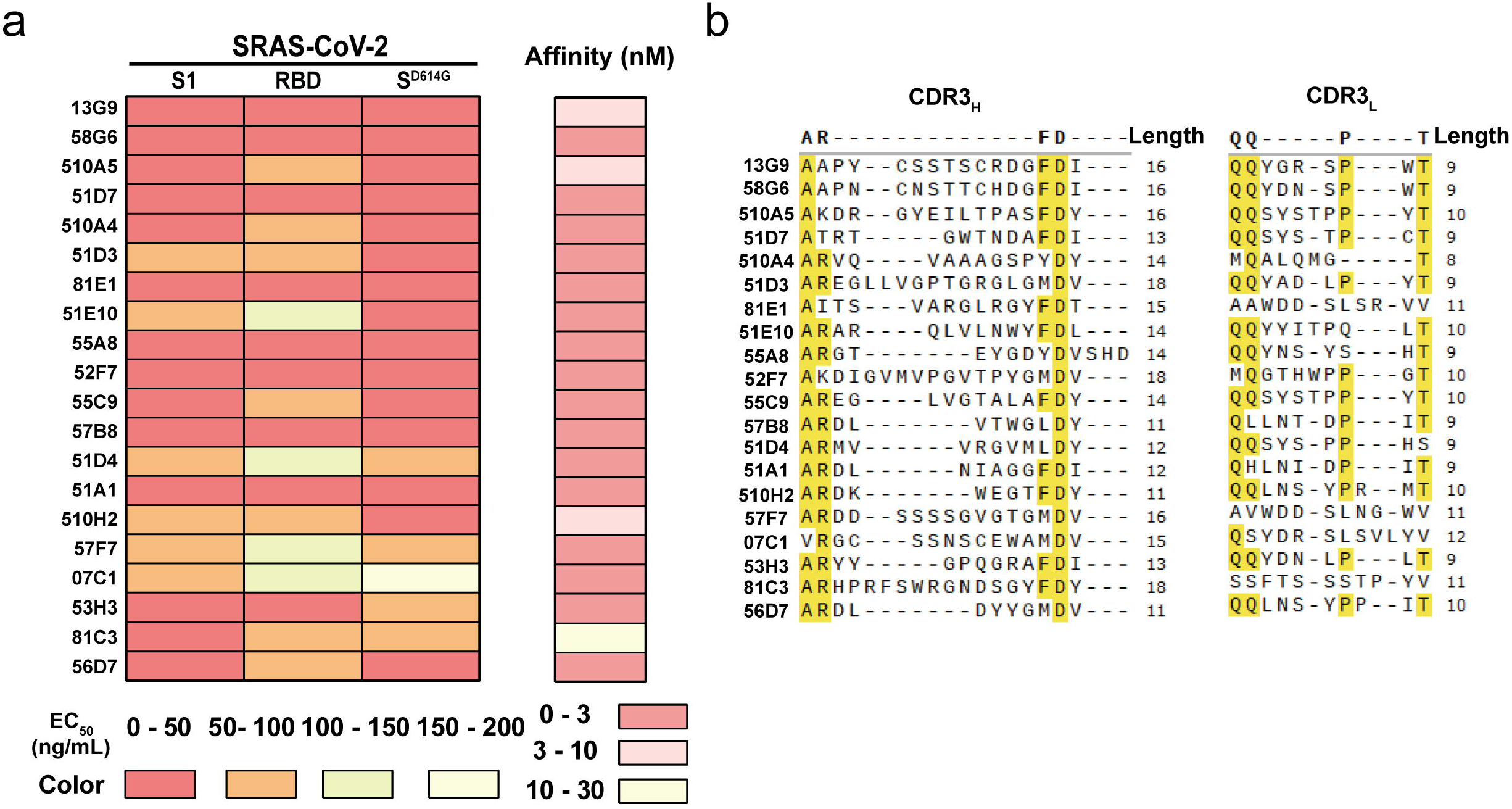
Characteristics of potential NAbs to SARS-CoV-2 S-RBD. (a) The heatmap of the selected NAbs binding to either SARS-CoV-2 S1, SARS-CoV-2 S-RBD or SARS-CoV-2 SD614G (left) and the affinity to SARS-CoV-2 S-RBD (right). The NAbs are ranked by their neutralizing potency to authentic SARS-CoV-2. The EC50 values of NAbs are visualized for binding. Data are representative of at least 2 independent experiments performed in technical duplicate. (b) The CDR3H (left) and CDR3L (right) sequences of the top 20 NAbs were aligned and the same amino acid residues were highlighted in yellow. The lengths of CDR3H and CDR3L sequence were labelled beside.

### Sequences of the potential NAbs to SARS-CoV-2 S-RBD

We further analyzed the gene sequence information of these potential NAbs, and observed diverse sequence pairs of the heavy- and light-chain variable regions (Extended Data Fig. 3). Several potential NAbs were preferentially transcribed from VH3-66 for the heavy chain. And VL1-9 and VL1-39 accounted for the light chain of about one third of these Abs (Extended Data Fig. 3). Their complementarity determining region 3 (CDR3) of the heavy- and light-chains exhibited a high variety of amino acid (aa) residues, with lengths ranging from 11 to18 aa and 8 to 12 aa for the heavy- and light-chains, respectively (Fig. 1b). These analyses demonstrated the genetic diversity of these potential NAbs from unique clonotypes.

### The neutralizing capabilities of the potential NAbs

In order to confirm the neutralizing capabilities of these potential NAbs, we performed virus neutralizing tests, quantified by the luminescence assay with pseudovirus bearing SARS-CoV-2 S and S^614G^, and by RT-qPCR with authentic SARS-CoV-2. Among these potential NAbs, the top 4 Abs (13G9, 58G6, 510A5 and 51D7) blocked authentic SARS-CoV-2 more efficiently, with the IC50 value of 0.007, 0.01, 0.011 and 0.039 μg/mL, respectively (Fig. 2a, Table 1). The similar neutralizing capabilities were observed in SARS-CoV-2 pseudovirus assays (Fig. 2b, Table 1). Two of them (58G6 and 510A5) showed consistent results with our previous findings^6^. The other 16 potential NAbs blocked authentic SARS-CoV-2 with the IC_50_ ranging from 0.043 to 0.153 μg/mL, and they neutralized SARS-CoV-2 pseudovirus with similar IC50 (Extended Data Fig. 4a, b, Table1). When we tested the inhibition effects of the top 4 NAbs against SARS-CoV-2 S^D614G^ pseudovirus, we found that these NAbs could also efficiently neutralize the pseudovirus, with IC_50_ value of 0.027, 0.018, 0.076 and 0.024 μg/mL, respectively (Fig. 2c, Table 1). We further confirmed that these NAbs blocked S-RBD-ACE2 interaction via SPR assay (Extended Data Fig. 5). Combining the results from previous sessions, as shown in Fig. 1, we found no correlation between the neutralizing activities and the binding affinity of the top 20 NAbs, suggesting that binding affinity, to a certain extent, was likely not a prerequisite for the neutralizing potency of NAbs (Extended Data Fig. 6). So far, at least 4 NAbs with high potency in neutralizing both authentic SARS-CoV-2 and SD^614G^ pseudovirus have been identified, which might be applied for potential interventions of the immediate epidemic.

**Fig. 2.**
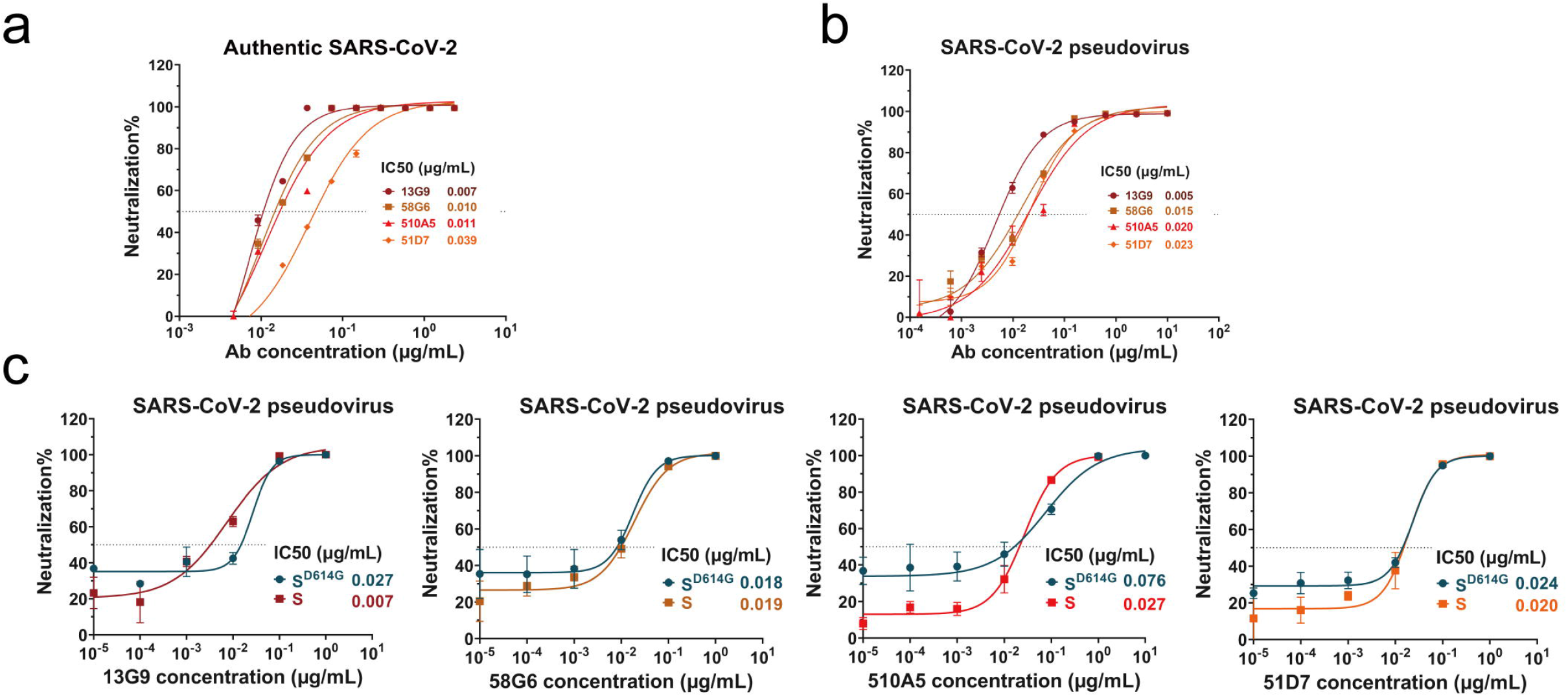
Assessment of the neutralizing capabilities of the NAbs against SARS-CoV-2 and its mutated S^D614G^. Neutralizing potency measured by the neutralization assay against authentic SARS-CoV-2 (a), SARS-CoV-2 pseudovirus (b) and its mutated type S^D614G^ (c). Data for each NAb were obtained from a representative neutralization experiment, with three replicates. Data are presented as mean ± SEM.

### The epitope mapping of the NAbs

To elucidate the potential mechanisms of NAb neutralizing SARS-CoV-2, we performed the epitope mapping for these NAbs via competitive ELISA assay. When we tested the above 20 NAbs and an additional 54 mAbs from our developed S-RBD-specific mAb reservoir, we found that they could be mainly divided into five clusters (Fig. 3a, Extended Data Fig. 7). Abs from each cluster exhibited relatively distinct preferences competing for the epitopes recognized by an NAb 13G9 (13G9e), a non-neutralizing Ab 81A11 (81A11e) and an NAb CR3022 to SARS-CoV (CR3022e) (Fig. 3a, Extended Data Fig. 7). All mAbs in the Cluster A competed with the most potent NAb 13G9, whereas each mAb in the Cluster C competed with 81A11 (Fig. 3a). Cluster B consisted of mAbs cross-reacted with 13G9e and 81A11e, the latter with a less content (Fig. 3a). Additionally, we found that all mAbs in the Cluster D were CR3022e-specific, all of them but one (07A10) strongly cross-reacted with both SARS-CoV S and S-RBD, indicating that CR3022e was relatively conserved between two coronaviruses (Fig. 3a, Extended Data Fig. 7, 8). The Cluster E did not recognize any of these three epitopes (Fig. 3a).

**Fig. 3.**
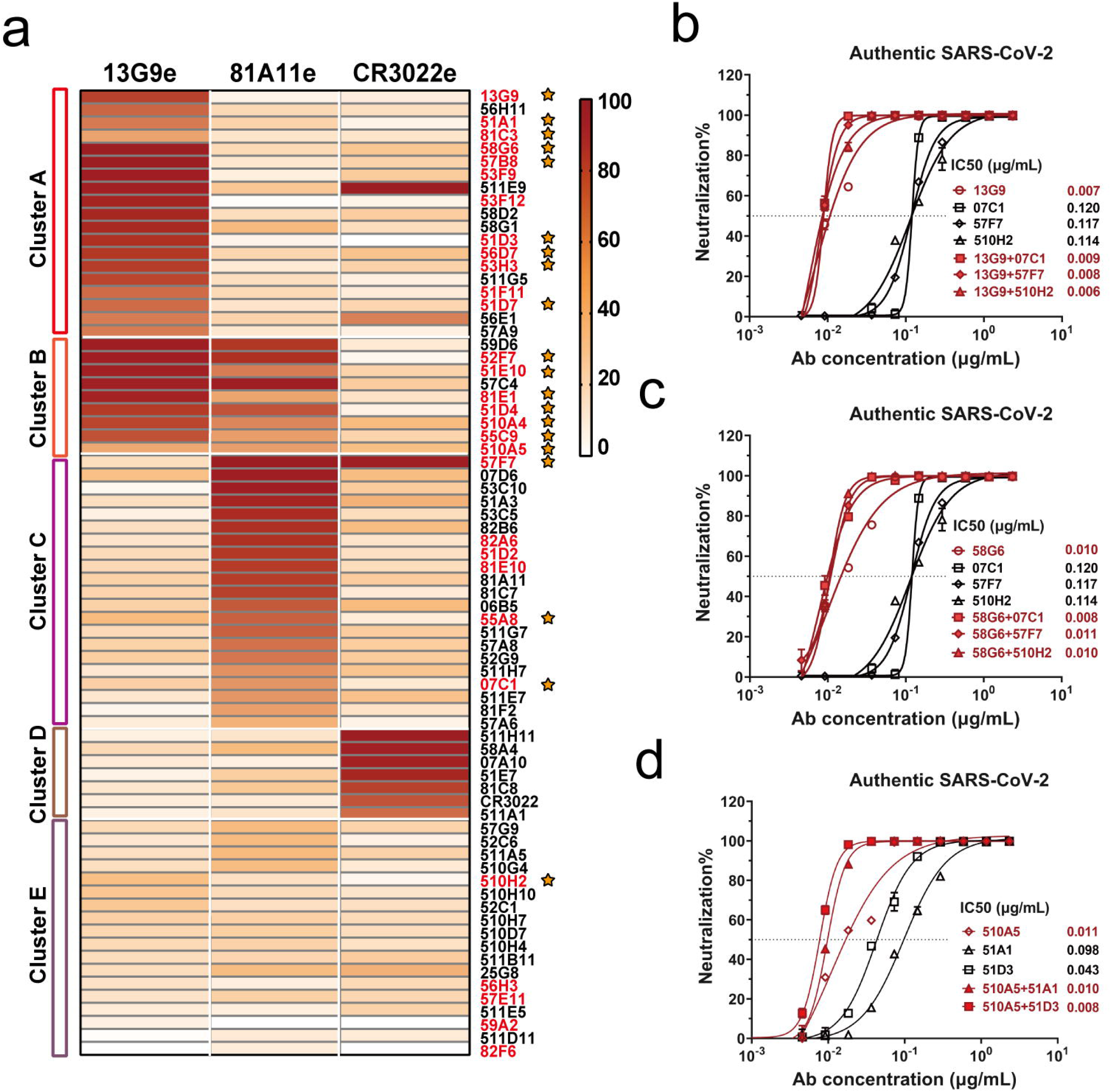
Epitope mapping of mAbs and the analysis of NAbs from different clusters. (a) Epitope mapping of purified mAbs targeting three independent epitopes (13G9e, 81A11e and CR3022e). The all NAbs detected by authentic SARS-CoV-2 CPE assay or SARS-CoV-2 pseudovirus neutralizing assay were labelled in red and the top 20 NAbs were indicated by orange stars. The combination effects of 13G9 (b) or 58G6 (c) with 07C1, 57F7 and 510H2, and 510A5 (d) with 51A1 and 51D3 against authentic SARS-CoV-2 were quantified by RT-qPCR, analyzing the efficiency of NAbs neutralization.

Interestingly, we found that 16 out of the 20 NAbs were grouped in Cluster A and Cluster B, inferring that at least 13G9e might be accounted for the binding of NAbs to S-RBD, whereas the epitope recognized by 81A11 (81A11e) was irrelevant (Fig. 3a). To further assess the interrelationships between the 20 NAbs in detail, we biotinylated them and performed the competitive ELISA assay. Majority of these NAbs exhibited similar competing patterns to that shown in Fig. 3a, while four NAbs recognized an independent epitope, suggesting that there might be at least two different epitopes responsible for the Nab and S-RBD interaction (Extended Data Fig. 9).

It has been hypothesized that applications with non-competitive pairs of NAbs might bring synergistic benefits for COVID-19 patients^20^. Hereby, we selected 13G9, 58G6 and 510A5 with the highest neutralizing potency from the Cluster A and B, and a few NAbs recognizing other epitopes, and tested their neutralizing activities in pairs against authentic SARS-CoV-2. As shown in Fig. 3b-d, synergy could be observed with 100% neutralization. However, the addition of NAbs from different clusters brought little advances to decrease the IC_50_ of potent NAbs, suggesting that our potent NAbs could be applied alone with sufficient neutralizing effect (Fig. 3b-d). Consistently, we found that a non-NAb 58A4 in the Cluster D, which targeted regions covering CR3022e, could efficiently block SARS-CoV with an IC50 of 0.033 μg/mL in the pseudovirus assay (Extended Data Fig. 10). The above results revealed that these NAbs with high potency shared at least one binding region, representing a key vulnerability site on the S-RBD, which played an important role blocking the entry of SARS-CoV-2 S or S^D614G^.

### A linear epitope within the vulnerability site on SARS-CoV-2 S-RBD

To further clarify the precise epitopes recognized by our potent NAbs, we investigated the antigenic profile of S-RBD targeted by our obtained NAbs. Based on the detection of linear epitopes by ELISA assay using 219 S-RBD-specific Abs, 15 mAbs were selected and further confirmed for their binding to denatured S-RBD, including 9 NAbs in the top 20 list (Extended Data Fig. 11). Subsequently, we designed and synthesized fifteen 20-mer peptides (RBD1 to RBD15), overlapping with 5 aa, to cover the entire RBD (S^319-541^) (Extended Data Fig. 12). Surprisingly, we found that the 10 of these 15 mAbs bound to RBD2, RBD9 and RBD13 simultaneously (Extended Data Fig. 13). Such results suggested that these 10 mAbs might target conformational epitopes bridging three independent linear regions, represented by RBD2, RBD9 and RBD13 (Extended Data Fig. 14). Furthermore, we observed that the mAbs with weak- or non-neutralizing abilities bound to all three peptides. Meanwhile, two of the most potent NAbs from the Cluster A, 58G6 and 51D3, exhibited similar binding preferences to RBD9, whereas they showed minimal interaction to RBD2 (Extended Data Fig. 13, 15). These results imply that RBD9 may represent a key region responsible for, consequently predicative of, NAb binding, whereas RBD2 may represent a region of steric interference that impedes effective interaction between RBD9-recognizing NAb and S-RBD.

Of note, the potent NAb 58G6 only bound to the RBD9 peptide, indicating that 58G6 interacted with a linear epitope on the S-RBD (Extended Data Fig. 15). To further identify essential aa residues in S-RBD accounted for 58G6 binding, we re-synthesized eight 20-mer peptides overlapping with 15 aa, namely RBD9-1 to RBD12-2 (Extended Data Fig. 12). Among these peptides, we found that 58G6 could bind to RBD9-1 in a dose-dependent manner, but not to RBD9-2, which shifted five aa residues from RBD9-1 (Fig. 4a, left). The structural studies from other groups have defined key aa residues for the S-RBD-ACE2 binding, specifically, G446, Y449, Y453, L455 and F456 located within the sequence represented by RBD9-1^12,18^. We individually replaced these aa residues with alanine (A) and found that the binding ability of 58G6 was markedly reduced with G446A or Y449A, and completely abolished with Y453A, L455A or F456A (Fig. 4a, right, Extended Data Fig. 16). Meanwhile, we assessed the interaction between RBD9-1 and ACE2, and similar results as those from 58G6 were obtained (Fig. 4b, Extended Data Fig. 16). Furthermore, the competition assay with S-RBD revealed that the linear epitope represented by RBD9-1 was an exclusive target of 58G6 (Fig. 4c). Additionally, we confirmed that 58G6 inhibited the interaction between ACE2 and RBD9-1, in a dose dependent manner (Fig. 4d). Collectively, these results confirmed that RBD9-1 represented a linear epitope for 58G6 (S^444-463^), as well as an essential binding region for SARS-CoV-2 S-RBD-ACE2 interaction (Extended Data Fig. 11). A very similar region of S-RBD (S^446-505^) was confirmed to be a critical steric site for ACE2 binding in an independent structural study^21^. In conclusion, we have discovered a linear epitope (S^444-463^) that is solely responsible for the high neutralizing potency of 58G6, and a key vulnerability site on SARS-CoV-2 S-RBD, which can provide important information for the NAb-based applications and vaccine design.

**Fig. 4.**
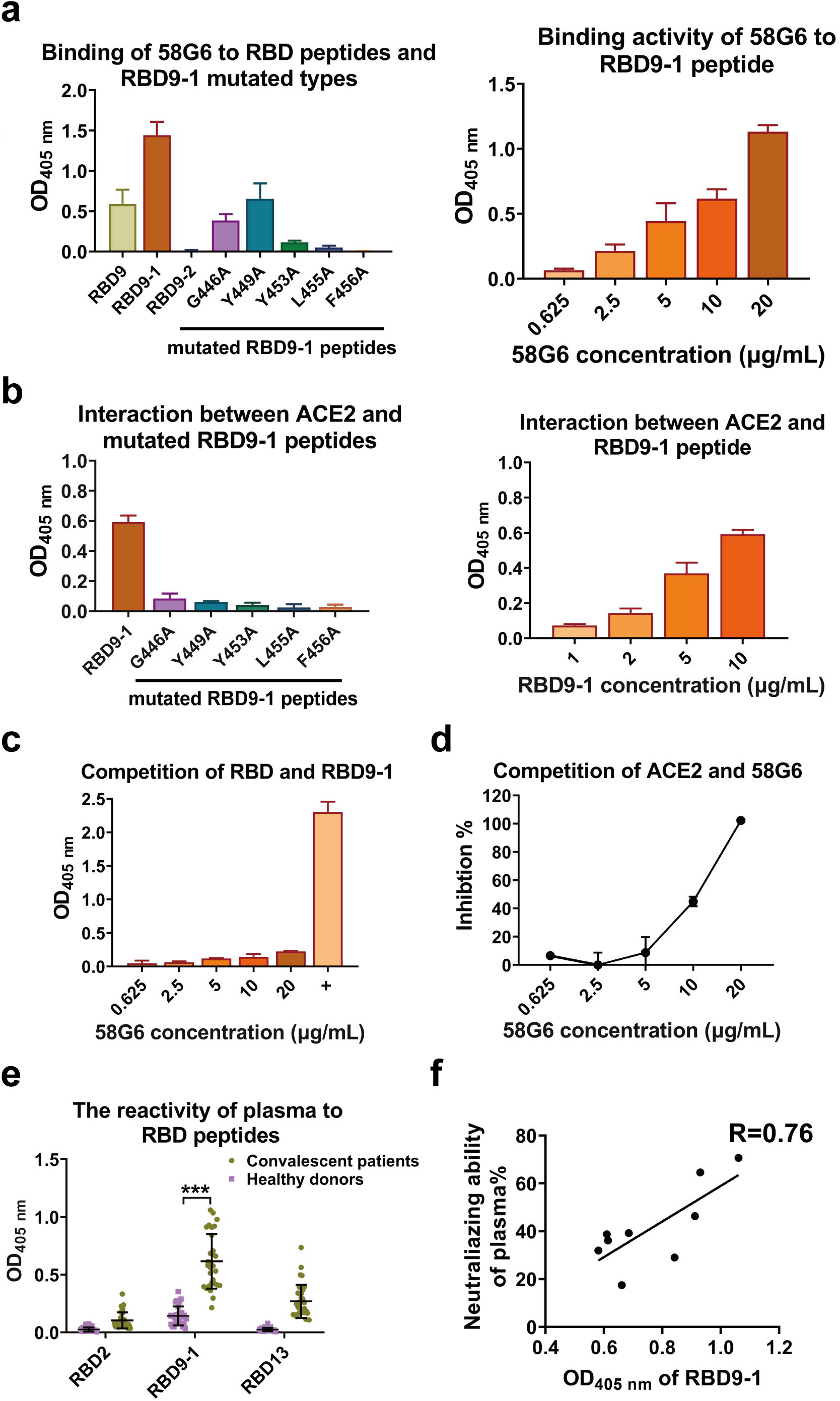
Linear epitope of 58G6 was a critical position in S-RBD-ACE2 binding region. (a) Peptides covering sequences in close proximity, RBD9, RBD9-1 and RBD9-2, were included to determine the specific region for 58G6 binding. The binding activity of 58G6 to the wild type (left) and the mutated (right) RBD9-1 peptides were tested by ELISA. (b) The interaction between ACE2 and wild type (right) and mutated (left) RBD9-1 peptides. (c) The ability of RBD9-1 blocking S-RBD and 58G6 interaction was analyzed by competitive ELISA. 1 μg/mL 58G6 was tested as positive control. (d) The ability of 58G6 blocking the interaction between RBD9-1 and ACE2, tested by competitive ELISA. (e) The reactivity of the plasma from convalescent patients and healthy donors with RBD2, RBD9-1 and RBD13 were assessed by ELISA. ***, p < 0.001. (f) The correlation between the neutralizing activity of the plasma from convalescent patients and the corresponding RBD9-1 binding activity, n=9.

Finally, to test whether the linear epitope could elicit a humoral immune response corresponding to that from SARS-CoV-2 in COVID-19 patients, we examined the interaction of RBD2, RBD9-1 or RBD13 peptide with convalescent plasma. RBD9-1 could react with a significantly larger amount of IgGs compared with those from healthy donors, whereas RBD2 and RBD13 both exhibited lower reactivities (Fig. 4e). Also, we confirmed that the all patients’ plasma samples were capable of neutralizing SRAS-CoV-2 pseudovirus, and such neutralizing abilities varied across patients (Extended Data Fig. 17). Importantly, we found a significant positive correlation between the neutralizing capabilities of the patients’ plasma against SARS-CoV-2 pseudovirus and their binding activities to RBD9-1 (Fig. 4f). This correlation revealed that 58G6e might be a potential biomarker used to detect the NAbs induced by SARS-CoV-2 infection or vaccination.

## Discussion

In our previous study, at least 50% S-RBD specific mAbs derived from the memory B cells were found to be NAbs^6^. Here, we focused on the top 20 potential NAbs for their functional characterization and the neutralizing mechanism study, in detail. Among these NAbs, we evidenced that 13G9, 58G6, 510A5 and 51D7 exhibited the highest neutralizing potency against authentic SARS-CoV-2, pseudovirus and its variant S^D614G^, with no obvious preference, indicating that these NAbs might be applicable for targeting mutated genotypes of non-RBD of SARS-CoV-2 S. Due to the significance of S^D614G^ in the current pandemic, all 4 NAbs may exhibit fitness advantages for the prophylaxis and the treatment of SARS-CoV-2 S^D614G^ infection. In some studies, researchers proposed the antibody cocktail strategies for targeting multiple distinct epitopes and for preventing rapid mutational escape resulted from individual antibodies^20,22^. In our study, no apparent differences of IC_50_ were observed between potent NAbs (13G9, 58G6 or 510A5) applied individually and in combination with NAbs from other clusters. We deemed that these NAbs alone could sufficiently inhibit authentic SARS-CoV-2 entry.

So far, NAbs have only been reported to recognize conformational epitopes, which provide limited information for vaccine design and targeting therapy. Recent studies identified several linear epitopes via convalescent patients’ plasma or the immunized mice^21,23^. Very fortunately, we discovered a linear epitope on S-RBD (S^444-463^) that could be exclusively recognized by one of the most potent NAbs (58G6). It represented, at least one of, the vulnerable sites on the RBD for SARS-CoV-2 neutralization. Significantly, 58G6 belongs to the Cluster A, which contains a variety of NAbs that can recognize at least a portion of the same conformational epitope 13G9e. Therefore, we can reasonably speculate that the mechanism determining the neutralizing potency may be shared by these NAbs from Cluster A and Cluster B, which may at least rely on the linear epitope represented by RBD9-1 (58G6e). Competition assay has confirmed that our NAbs targeting S^444-463^ play crucial roles interfering with the RBD-ACE2 interaction, which determines the virus entry. Importantly, the convalescent patients’ plasma with neutralizing activity can bind to RBD9-1, implying that RBD9-1 might be used for serological diagnosis. The immune response to 58G6e, a candidate of immunogen, is currently being investigated with animal models, for the development of SARS-CoV-2 vaccines.

In this study, we have obtained several NAbs that yield a complete blockage of authentic SARS-CoV-2 and SASR-CoV-2 S^D614G^ pseudovirus. We are the first, as far as we know, to provide direct evidence of a linear epitope that can be recognized by a potent NAb against SARS-CoV-2 S-RBD. Our findings have illuminated potential mechanisms for the neutralizing activity of NAbs, which paves the way for possible safe and effective NAb treatment and vaccine design.

## Methods

### Antibodies

The human antibodies studied in this paper were isolated from 39 SARS-CoV-2 convalescent patient plasma in China. The original clinical studies to obtain blood samples after written informed consent were previously described^6^ and had been approved by the Ethics Board of ChongQing Medical University. The antibodies were isolated using flow sorting for isolation and cloning of single antigen-specific B cells and the antibody variable genes encoding monoclonal antibodies^6^.

### Recombinant antibody production and purification

For the construction of antibody expression Vectors, VH and VL 2nd PCR products were inserted separately into the linearized plasmids (pcDNA3.4) that encode constant regions of the heavy chains and light chains via a homologous recombination kit (Vazyme, C112). A pair of plasmids separately expressing the heavy- and light-chain of antibodies were transiently co-transfected into Expi293™ cells (Catalog No. A14528, ThermoFisher) with ExpiFectamine™ 293 Reagent. Then the cells were cultured in shaker incubator at 120 rpm and 8 % CO_2_ at 37 □. After 7 days, the supernatants with the secretion of antibodies were collected and captured by protein G Sepharose (GE Healthcare). The bound antibodies on the Sepharose were eluted and dialyzed into phosphate-buffered saline (PBS). The purified antibodies were used in following binding and neutralization analyses.

### Sequence analysis of antigen-specific mAb

IMGT/V-QUEST (http://www.imgt.org/IMGT_vquest/vquest) and IgBLAST (https://www.ncbi.nlm.nih.gov/igblast/), MIXCR (https://mixcr.readthedocs.io/en/master/) and VDJtools (https://vdjtools-doc.readthedocs.io/en/master/overlap.html) tools were used to do the variable region analysis and annotation for each antibody clone.

### MAb quantification

Purified mAbs quantification was performed by UV spectrophotometry using a NanoDrop spectrophotometer and accounting for the extinction coefficient of human IgG.

### The Antibody binding kinetics and the competition with ACE2 measured by SPR

The affinity between the NAbs and SARS-CoV-2 S-RBD were measured using the Biacore X100 platform at room temperature (RT). A CM5 chip (GE Healthcare) was linked with anti-human IgG-Fc antibody to capture about 9000 response units of the NAbs. The gradient concentrations of SARS-CoV-2 S-RBD were prepared (2-fold dilutions, from 50 nM to 0.78125 nM) using HBS-EP^+^ Buffer (0.01 M HEPES, 0.15 M NaCl, 0.003 M EDTA and 0.05% (v/v) Surfactant P20, pH 7.4), and sequentially injected into the chip and monitored for the binding kinetics. After the final reading, the sensor surface of the chip was regenerated with 3 M MgCl2 (GE) before the measurement of the next NAb. The affinity of the NAbs was calculated with Biacore X100 Evaluation Software (Version:2.0.2) using 1:1 binding fit model.

To determine competition with the human ACE2 peptidase domain, SARS-CoV-2 S-RBD was coated on a CM5 sensor chip via amine group for a final RU around 250. Antibodies (20 μg/mL) were injected onto the chip until binding steady-state was reached. ACE2 (20 μg/mL) was then injected for 60 seconds. Blocking efficacy was determined by comparison of response units with and without prior antibody incubation.

### ELISA

The tests of Nabs reactivities against antigen were performed as previous reports^24^. Briefly, 2 μg/mL recombinant S1 or S-RBD of SARS-CoV-2 or SARS-CoV (Sino Biological, Beijing, China) were coated on 384-well plates (Corning Costar), and stored at 4 °C overnight. The next day, the plates were first blocked with the blocking buffer (PBS with 5% BSA) at 37 °C for 1 h, and then added with serially diluted mAbs (10-fold dilutions, from 10 μg/mL to 10 pg/mL). After 30 min incubation at 37 °C, the plates were washed 5 times and incubated with goat anti-human IgG (H+L) antibody conjugated with ALP (Thermo Fisher, a18808, 1:5000) for 30 min at 37 °C. For the quantification of bound IgG, PNPP (Thermo Fisher) was added at 1 mg/mL and the absorbance at 405 nm was measured by the MultiSkan GO fluoro-microplate reader (Thermo Fisher).

### Competitive ELISA

For the competitive ELISA used in epitope mapping of mAbs, 2 μg/mL recombinant S-RBD-his (Sino Biological, Beijing, China) was added in 384-well plates and incubated at 4 °C overnight. 50 μg/mL mAbs per well were added. The plates were incubated at 37 °C for 1 h and then washed. Biotinylation of mAbs (the top 20 Nabs and 13G9, 81A11, previously reported SARS-CoV CR3022^25^) were performed using the EZ-link NHS-PEO Solid Phase Biotinylation Kit (Pierce) according to the manufacturer’s protocol and purified using MINI Dialysis Unit (ThermoFisher, 69576). 500 ng/mL biotinylated mAbs were added to each well, and the plates were incubated at 37 °C for 1 h. ALP-conjugated Streptavidin (Mabtech, Sweden, 3310-10) was added at 1:1000, followed by an incubation of 30 min at 37 °C. Quantification step was the same as described above.

### peptide ELISA

Peptide ELISA was performed with synthesized peptides overlapping with 5 amino acids (Genescripts, Wuhan, China). These peptides were tethered by N-terminal biotinylated linker peptides (biotin-ahx), except for the first peptide at the N-terminus, whose biotin was linked to the C terminus instead. The RBD9-1 amino acid residues were selected and mutated to alanine according to the previous structural reports^12^ and synthesized by Genescripts (Wuhan, China). 50 μL synthesized peptide was added to the streptavidin-coated 384-well plate in duplets to make a final concentration of 5 μg/mL. The plates were incubated for 2 hrs at RT. After washing, the plates were blocked with Protein-Free Blocking Buffer (Pierce, USA, 37573) at RT for 1 h and incubated with 10 μg/mL testing mAbs or COVID-19 convalescent patients’ plasma (1:100) at RT for another 1 h. Reacted mAbs were detected using ALP-conjugated Goat F(ab’)2 Anti-Human (IgG (Fab’)2) secondary antibody (Abcam, ab98532, 1:2000) for 30 min at RT, followed with quantification detection.

For the ACE2 competitive peptide ELISA, 5 μg/mL synthesized RBD9-1 was immobilized on the streptavidin-coated 384-well plate at RT for 2 hrs. After washing with Protein-Free Blocking Buffer, the plates were blocked with this Blocking Buffer. Next, serial diluted 58G6 (20-0.625 μg/mL) in 50 μL of the Blocking Buffer were added into plate and the plates were incubated at RT for 1 h. Then, the plate incubated with 2 μg/mL ACE2 at RT for another 1 h. The ELISA plates were washed 4 times by Blocking Buffer and 50 μL Goat F(ab’)2 Anti-Human (IgG (Fab’)2) secondary antibody conjugated with ALP (Abcam, ab98532, 1:2000) was incubated with the plate at RT for 30 min. The plate was washed and followed with quantification detection.

For the RBD competitive peptide ELISA, 5 μg/mL synthesized RBD9-1 was coated on the plate at RT for 2 hrs. Then, the plate was blocked with Protein-Free Blocking Buffer. 50 μL serial diluted 58G6 (20-0.625 μg/mL) was added into each well in the plates and the plates were incubated at RT for 1 h. Next, 2 μg/mL RBD-mFC (Sino Biological, Beijing, China) was added to each well, and the plates were incubated at RT for 1 h. The ELISA plates were washed 4 times by Blocking Buffer and ALP-conjugated Goat Anti-Mouse IgG Fc (Abcam, ab98710, 1:1000) in 50 μL Blocking Buffer was added. Quantification step was the same as described above.

### Production of S protein pseudovirus

pVSVG expressing SARS-CoV-2 spike was constructed as previously described^26^. The packaging plasmid (VSV-G pseudotyped ΔG-luciferase) encoding either SARS-CoV S, SARS-CoV-2 S and SARS-CoV-2 S^D614G^ was generated. HEK293T cells were grown to 80% confluency before transfection with VSV-G pseudotyped ΔG-luciferase, pWPXL and pSPAX2. Cells were cultured overnight at 37 □°C with 5% CO_2_. DMEM supplemented with 5% fetal bovine serum and 100 I.U./mL of penicillin and 100 μg/mL of streptomycin were added to the inoculated cells, which were cultured overnight for 72 h. The supernatant was harvested, filtered by 0.45 μm filter and centrifugated at 300 g for 10 min to collect the supernatant and then aliquoted and storied at −80°C.

### Pseudovirus neutralization assay

50 μL serially diluted mAbs or convalescent patients’ plasma (1:1000) were incubated with the same volume of the HEK293T cell supernatants containing pseudovirus for 1 h at 37 °C. These pseudovirus-antibody mixtures were added to ACE2 expressing HEK293T cells (HEK293T/ACE2). After 72 hrs, the luciferase activities of infected HEK293T/ACE2 cells were detected by the Bright-Luciferase Reporter Assay System (Promega, E2650). The half-maximal inhibitory concentrations (IC50) of the evaluated mAbs were tested by the Varioskan LUX Microplate Spectrophotometer (Thermo Fisher), and calculated by a four-parameter logistic regression using GraphPad Prism 8.0.

### Authentic SARS-CoV-2 neutralization assay

This assay was performed in a biosafety level 3 laboratory of Fudan University. Serially diluted mAbs were incubated with authentic SARS-CoV-2 (nCoV-SH01, GenBank: MT121215.1, 100 TCID50) for 1 h at 37 °C. After the incubation, the mixtures were then transferred into 96-well plates, which were seeded with Vero E6 cells. The plates were kept at 37 °C for 48 hrs and each well was visually assessed for the cytopathic effect (CPE). And the supernatant viral RNA load of each well was quantified by RT-qPCR.

For RT-qPCR, the viral RNA was extracted from the collected supernatant using Trizol LS (Invitrogen) and used as templates for the RT-qPCR analysis by Verso 1-Step RT-qPCR Kit (Thermo Scientific) following the manufacturer’s instructions. PCR primers targeting SARS-CoV-2 N gene (nt 608-706) were as followed, forward: 5’-GGGGAACTTCTCCTGCTAGAAT-3’, and reverse: 5’-CAGACATTTTGCTCTCAAGCTG-3’. RT-qPCR was performed using the LightCycler 480 II PCR System (Roche) with the following program: 50°C 15 min; 95°C 15 min; 40 cycles of 95°C 15 sec, 50°C 30 sec, 72°C 30 sec.

### Western blot analysis

The recombinant S-RBD protein was mixed with 5× loading buffer (Beyotime, Shanghai, China) and denatured for 5 min at 100°C. Denatured proteins (200 ng) were subjected to electrophoresis with 10 % SDS-polyacrylamide gel and then transferred to PVDF membranes. After blocking by skim milk (Biofroxx), the membranes were incubated at 4 °C overnight, with the purified mAbs as primary Abs. The next days, the membranes were washed with TBST and incubated with HRP-conjugated Goat-anti-human Fc antibody (Abcam, ab99759, 1:10000) for 1 h at RT. The membranes were examined on Bio-rad ChemiDoc Imaging System (Bio-rad).

### Data analysis

Data are shown as mean ± SEM. Two-group comparisons were performed by Student’s t-test. The difference was considered significant if p < 0.05.

## Data Availability

Materials reported in this study will be made available but may require execution of a Materials Transfer Agreement. Further information and requests for resources and reagents should be directed to and will be fulfilled by the corresponding author Aishun Jin.

## Ethics Statement

The project ‘The application of antibody tests patients infected with SARS-CoV-2’ was approved by the Ethics Board of ChongQing Medical University. Informed consents were obtained from all participants.

## Acknowledgments

We thank health donors from Chongqing Medical University. We thank M. Zhu from Department of laboratory, The Third Affiliated Hospital of Chongqing Medical University. This study was supported by Chongqing Medical University fund (X4457) with the donation from Mr. Yuling Feng.

## Author contributions

A.J. and A.H. conceived and designed the study. F.G. and K.D. were responsible for recruiting patients and collecting the plasma and peripheral blood mononuclear cell., F.L. and H.J. were responsible for antibody production and purification. J.W., K.W., J.H., S.L., N.T., G.Z. and Q.G. conducted the pseudovirus neutralization assays and Y.X., C.G., Y.W., W.X., X.C., D.Q. and Z.Y. performed authentic SARS-CoV-2 neutralization assays. S.L. and Y.H. played an import role in data analysis of neutralizing Abs sequences. T.L., Y.W., Y.L., S.S. Q.C., F.G., M.S. and W.W. performed ELISA, competitive ELISA and peptide ELISA. X.H., C.H., R.W. and S.M. were responsible for SPR assay for the affinity of these NAbs and competition of these NAbs with ACE2. L.L., C.H., Y.N. and L.D. generated figures and tables, and take responsibility for the integrity and accuracy of the data presentation. A.J., T.L. and Y.X. wrote the manuscript.

## Competing interests

Patent has been filed for some of the antibodies presented here.

## Additional Information

Correspondence and requests for materials should be addressed to A.J.

## Notes

### Competing Interest Statement

The authors have declared no competing interest.

## References

1 Guan, W. J. et al. Clinical Characteristics of Coronavirus Disease 2019 in China. The New England journal of medicine 382, 1708–1720, doi:10.1056/NEJMoa2002032 (2020).

2 Lu, R. et al. Genomic characterisation and epidemiology of 2019 novel coronavirus: implications for virus origins and receptor binding. Lancet (London, England) 395, 565–574, doi:10.1016/s0140-6736(20)30251-8 (2020).

3 Zhou, P et al. A pneumonia outbreak associated with a new coronavirus of probable bat origin. Nature 579, 270–273, doi:10.1038/s41586-020-2012-7 (2020).

4 Korber, B. et al. Tracking Changes in SARS-CoV-2 Spike: Evidence that D614G Increases Infectivity of the COVID-19 Virus. Cell, doi:10.1016/j.cell.2020.06.043 (2020).

5 Toyoshima, Y., Nemoto, K., Matsumoto, S., Nakamura, Y. & Kiyotani, K. SARS-CoV-2 genomic variations associated with mortality rate of COVID-19. Journal of human genetics, 1–8, doi:10.1038/s10038-020-0808-9 (2020).

6 Han, X. et al. A rapid and efficient screening system for neutralizing antibodies and its application for the discovery of potent neutralizing antibodies to SARS-CoV-2 S-RBD. bioRxiv, 2020.2008.2019.253369, doi:10.1101/2020.08.19.253369 (2020).

7 Zhang, L. et al. The D614G mutation in the SARS-CoV-2 spike protein reduces S1 shedding and increases infectivity. bioRxiv, 2020.2006.2012.148726, doi:10.1101/2020.06.12.148726 (2020).

8 Hu, J. et al. The D614G mutation of SARS-CoV-2 spike protein enhances viral infectivity and decreases neutralization sensitivity to individual convalescent sera. bioRxiv, 2020.2006.2020.161323, doi:10.1101/2020.06.20.161323 (2020).

9 Hoffmann, M. et al. SARS-CoV-2 Cell Entry Depends on ACE2 and TMPRSS2 and Is Blocked by a Clinically Proven Protease Inhibitor. Cell 181, 271–280.e278, doi:10.1016/j.cell.2020.02.052 (2020).

10 Letko, M., Marzi, A. & Munster, V Functional assessment of cell entry and receptor usage for SARS-CoV-2 and other lineage B betacoronaviruses. Nature microbiology 5, 562–569, doi:10.1038/s41564-020-0688-y (2020).

11 Yan, R. et al. Structural basis for the recognition of SARS-CoV-2 by full-length human ACE2. Science (New York, N.Y.) 367, 1444–1448, doi: 10.1126/science.abb2762 (2020).

12 Lan, J. et al. Structure of the SARS-CoV-2 spike receptor-binding domain bound to the ACE2 receptor. Nature 581, 215–220, doi:10.1038/s41586-020-2180-5 (2020).

13 Malonis, R. J., Lai, J. R. & Vergnolle, O. Peptide-Based Vaccines: Current Progress and Future Challenges. Chemical reviews 120, 3210–3229, doi: 10.1021/acs.chemrev.9b00472 (2020).

14 Long, Q. X. et al. Antibody responses to SARS-CoV-2 in patients with COVID-19. Nature medicine 26, 845–848, doi:10.1038/s41591-020-0897-1 (2020).

15 Cao, Y. et al. Potent Neutralizing Antibodies against SARS-CoV-2 Identified by High-Throughput Single-Cell Sequencing of Convalescent Patients’ B Cells. Cell 182, 73–84.e16, doi:10.1016/j.cell.2020.05.025 (2020).

16 Ju, B. et al. Human neutralizing antibodies elicited by SARS-CoV-2 infection. Nature 584, 115–119, doi:10.1038/s41586-020-2380-z (2020).

17 Liu, L. et al. Potent neutralizing antibodies against multiple epitopes on SARS-CoV-2 spike. Nature, doi:10.1038/s41586-020-2571-7 (2020).

18 Shi, R. et al. A human neutralizing antibody targets the receptor-binding site of SARS-CoV-2. Nature 584, 120–124, doi:10.1038/s41586-020-2381-y (2020).

19 Robbiani, D. F. et al. Convergent antibody responses to SARS-CoV-2 in convalescent individuals. Nature 584, 437–442, doi:10.1038/s41586-020-2456-9 (2020).

20 Baum, A. et al. Antibody cocktail to SARS-CoV-2 spike protein prevents rapid mutational escape seen with individual antibodies. Science (New York, N.Y.) 369, 1014–1018, doi:10.1126/science.abd0831 (2020).

21 Zhang, B. Z. et al. Mining of epitopes on spike protein of SARS-CoV-2 from COVID-19 patients. Cell research 30, 702–704, doi:10.1038/s41422-020-0366-x (2020).

22 Wu, Y et al. A noncompeting pair of human neutralizing antibodies block COVID-19 virus binding to its receptor ACE2. Science (New York, N.Y.) 368, 1274–1278, doi:10.1126/science.abc2241 (2020).

23 Lu, S. et al. The immunodominant and neutralization linear epitopes for SARS-CoV-2. bioRxiv, 2020.2008.2027.267716, doi:10.1101/2020.08.27.267716 (2020).

## References

24 Jin, A. et al. A rapid and efficient single-cell manipulation method for screening antigen-specific antibody-secreting cells from human peripheral blood. Nature medicine 15, 1088–1092, doi:10.1038/nm.1966 (2009).

25 ter Meulen, J. et al. Human monoclonal antibody combination against SARS coronavirus: synergy and coverage of escape mutants. PLoS medicine 3, e237, doi:10.1371/journal.pmed.0030237 (2006).

26 Ou, X. et al. Characterization of spike glycoprotein of SARS-CoV-2 on virus entry and its immune cross-reactivity with SARS-CoV Nature Communications 11, 1620, doi:10.1038/s41467-020-15562-9 (2020).

