## Supplemental Figure and Table for "A key linear epitope for a potent neutralizing antibody to SARS-CoV-2 S-RBD"

Includes:

Extended Data Fig. 1-17

Extended Data Table 1

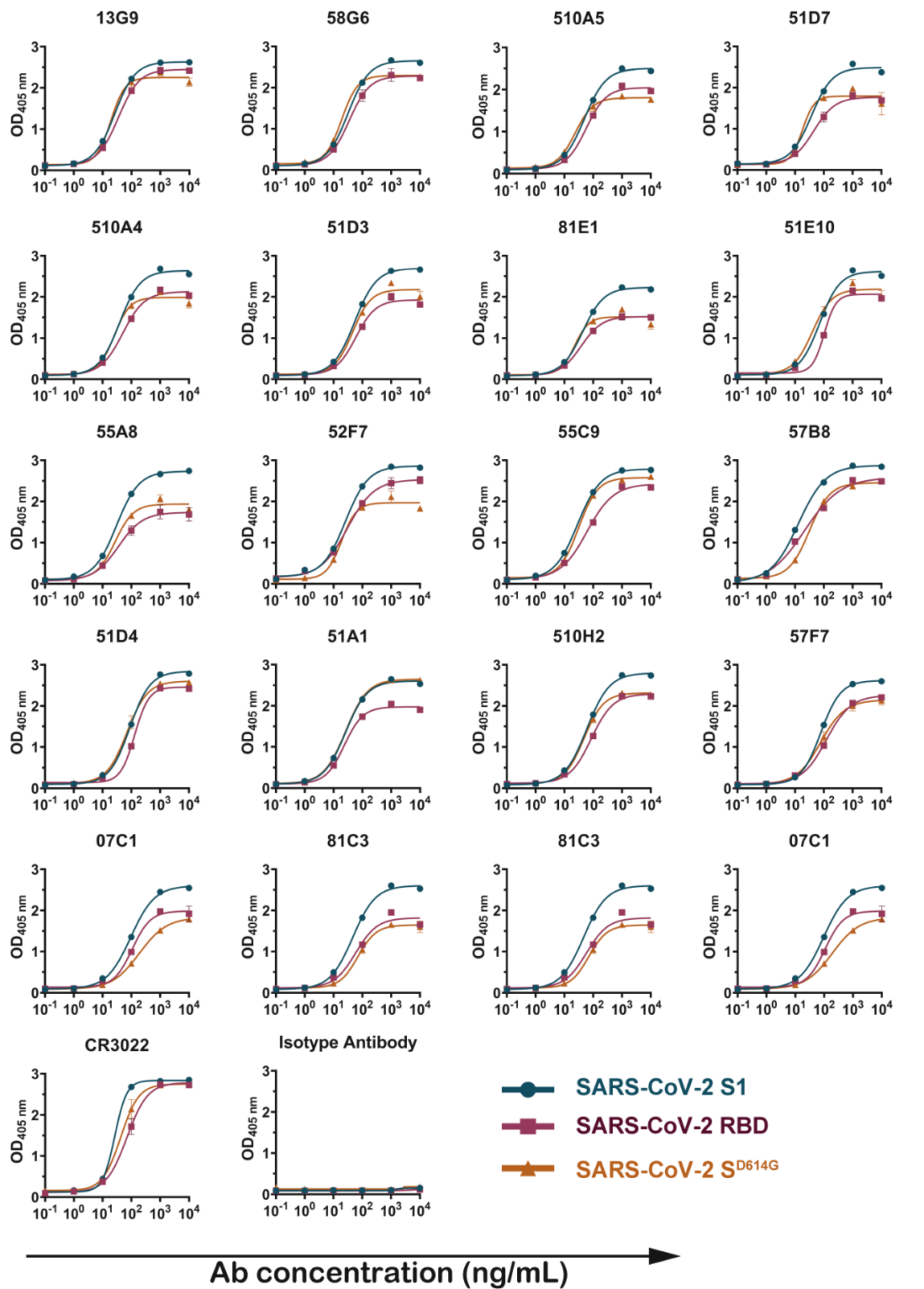
**Extended Data Fig. 1 The binding of the selected S-RBD specific NAbs to SARS-CoV-2 S1, SARS-CoV-2 S-RBD and SARS-CoV-2 S^D614G^.** The binding of these S-RBD specific NAbs to different targets were tested with various concentrations of NAbs by ELISA. The mAb CR3022 that cross-reacted with SARS-CoV S-RBD were tested as positive control and the HBV antibody were used as negative control. Data are representatives of at least 2 independent experiments performed in technical duplicate. The mean ± SEM of duplicates are shown.

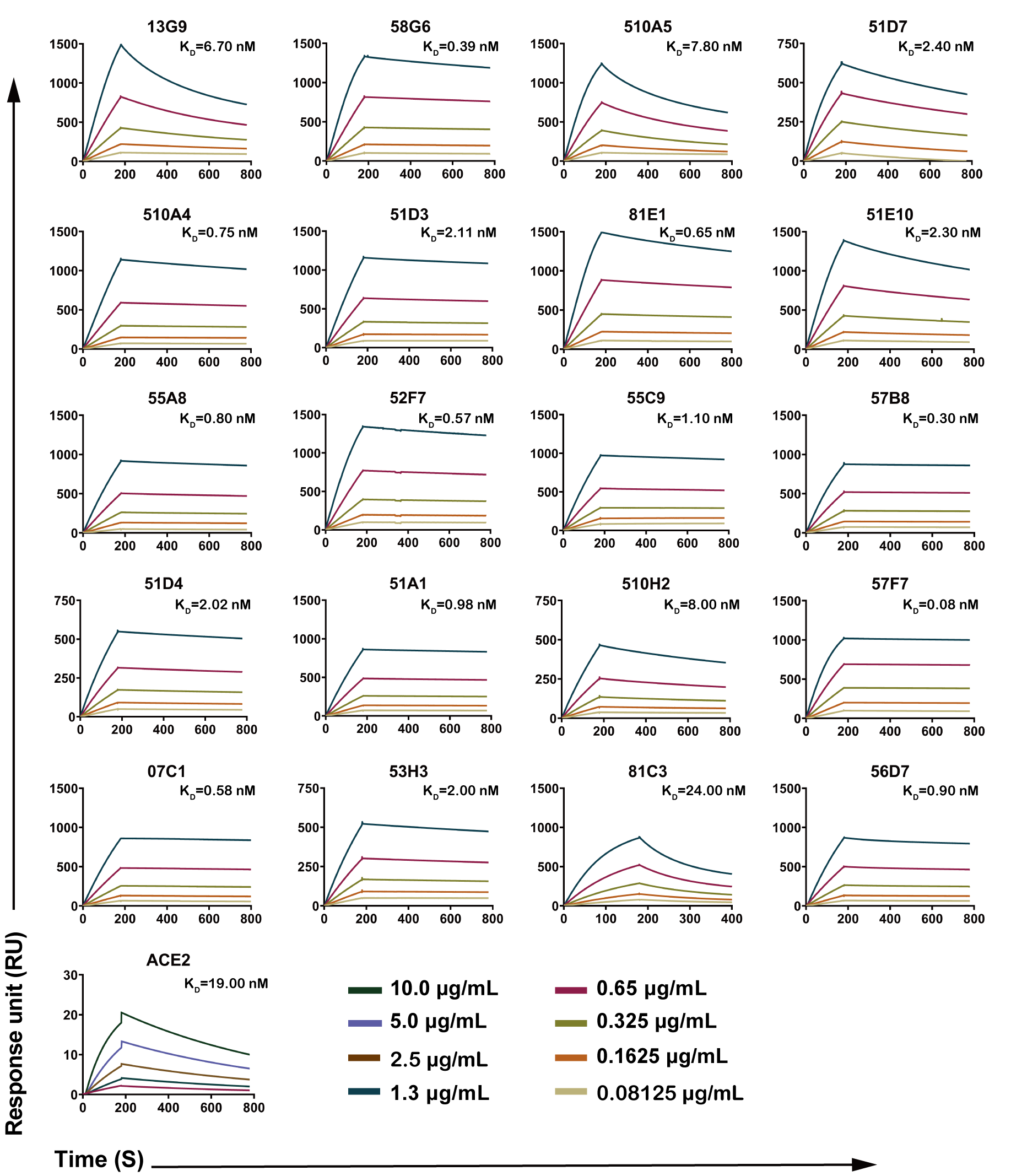

**Extended Data Fig. 2 The binding kinetics of the top 20 S-RBD specific NAbs with SARS-CoV-2 S-RBD.** The purified S-RBD NAbs were coated on the CM5 sensor chip followed by the injection of various concentrations of soluble SARS-CoV-2 S-RBD protein. For the assessment of ACE2 affinity, S-RBD was coated on the CM5 sensor chip, with subsequent injections of various concentrations of ACE2 to the coated chip. Data are representatives of at least 2 independent experiments.

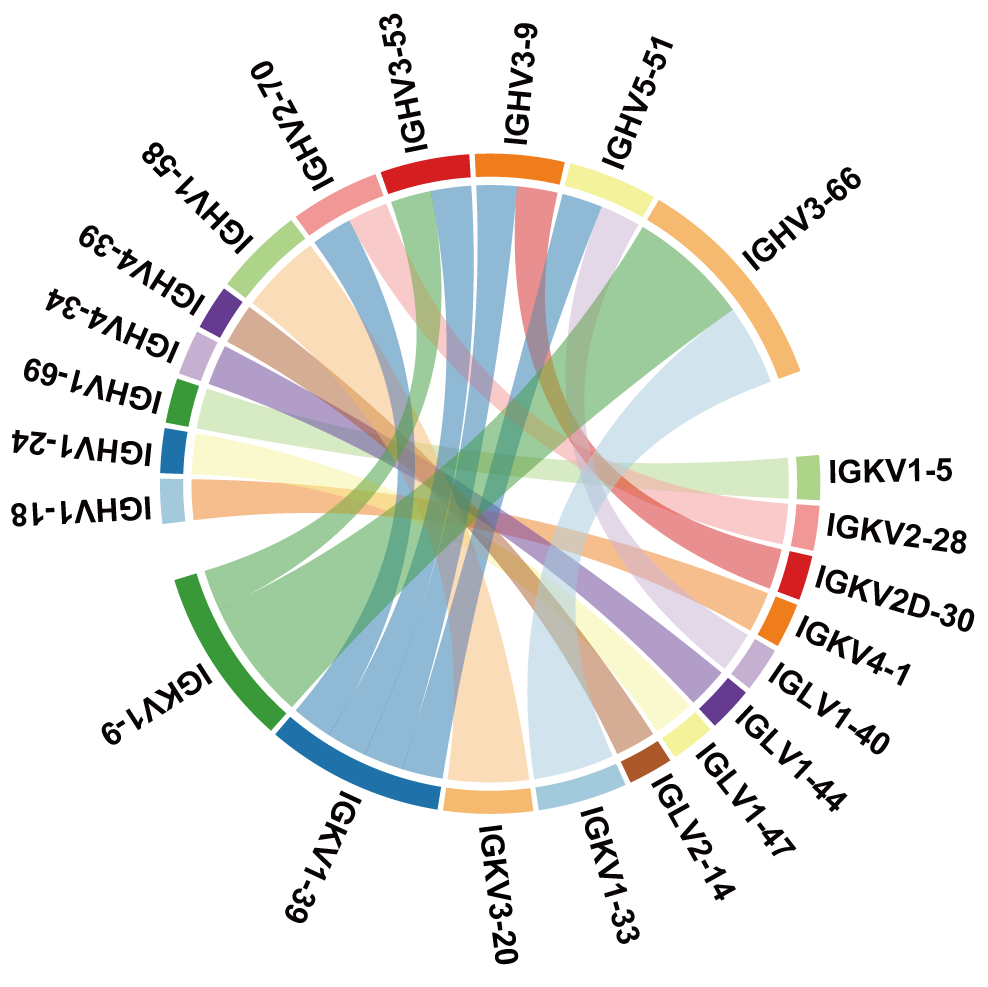

**Extended Data Fig. 3 The paired cluster information of the top 20 NAbs.** Both full-length heavy- and light-chain variable regions of the top 20 NAbs were paired. The bands between two clusters represent the pairing of the heavy-chain and light-chain variable regions. The widths of bands represent the frequencies of pairing.

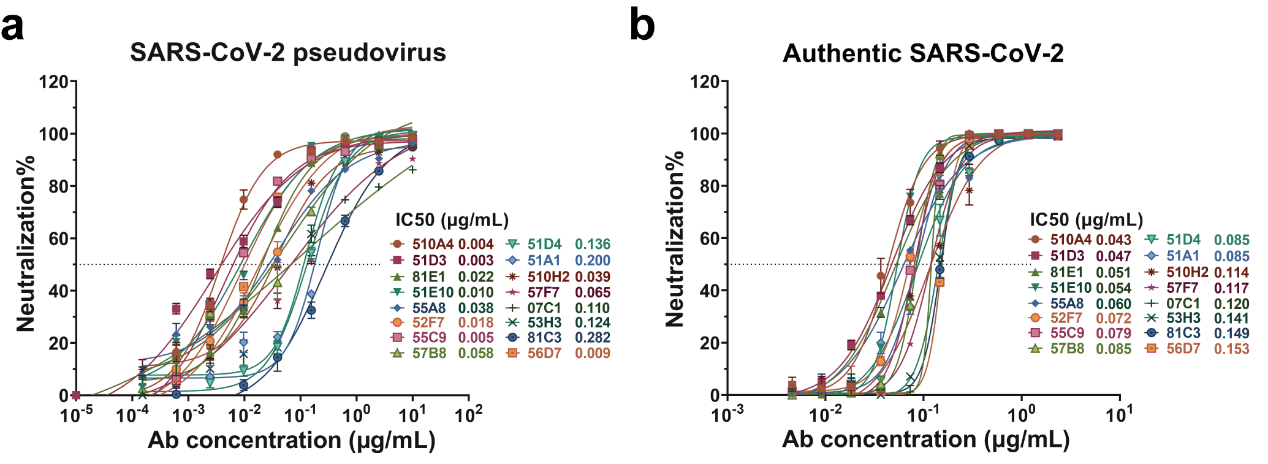

**Extended Data Fig. 4 The neutralizing capabilities of additional NAbs to SARS-CoV-2 pseudovirus or the authentic virus.** Neutralizing capabilities measured by the neutralization assay against SARS-CoV-2 pseudovirus (a) or the authentic virus (b). Data shown for each NAb were obtained from a representative neutralization experiment, with three replicates. Data are presented as mean ± SEM.

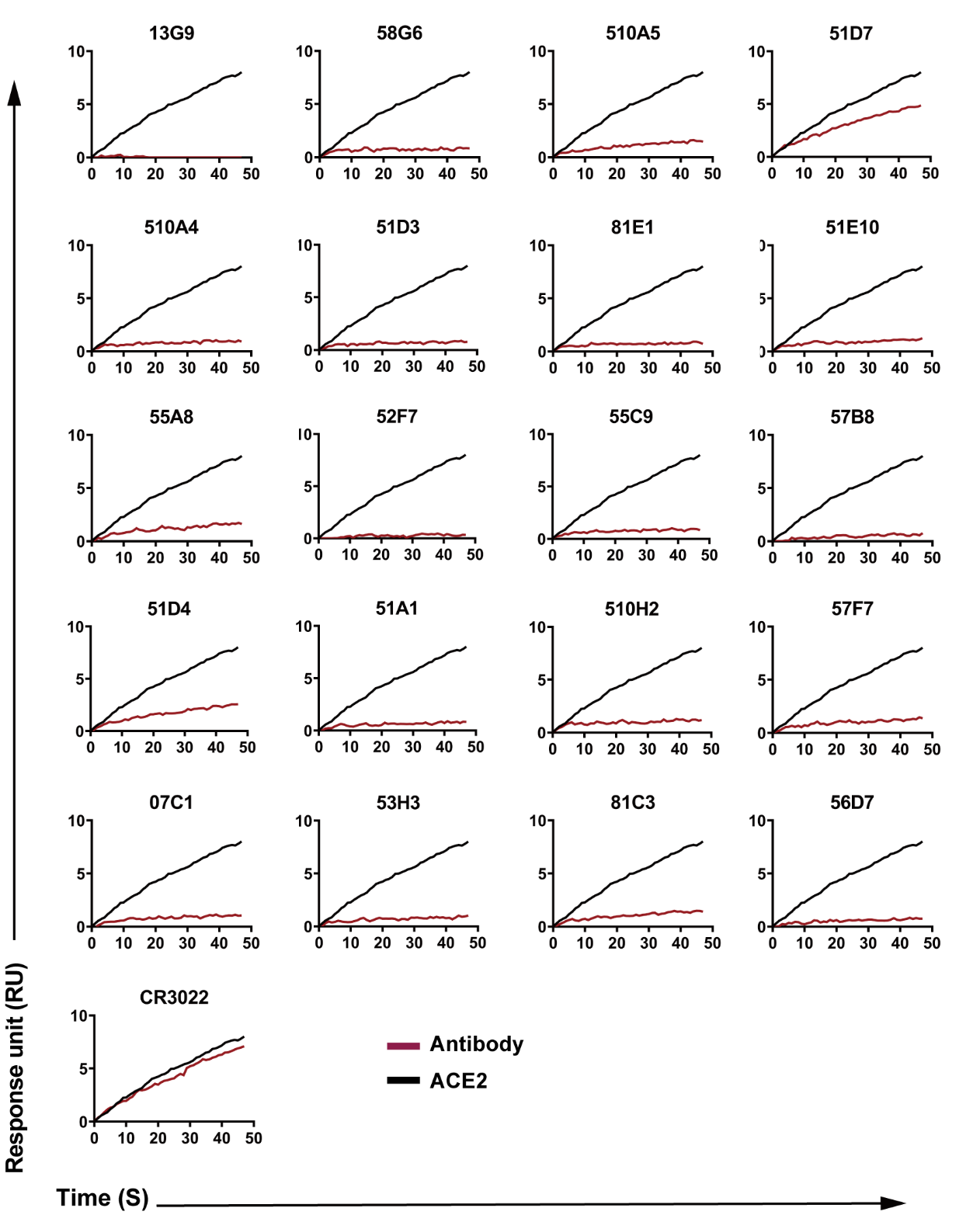

**Extended Data Fig. 5 The competition between S-RBD and ACE2.** The purified soluble SARS-CoV-2 S-RBD protein was covalently coated onto a CM5 sensor chip, followed by injections of individual NAbs at a concentration of 20 μg/mL. The competition capacity of each antibody is indicated by the level of reduction in response to ACE2, comparing to the corresponding level without prior antibody incubation. The SARS-CoV S-RBD mAb CR3022 was tested as the negative control.

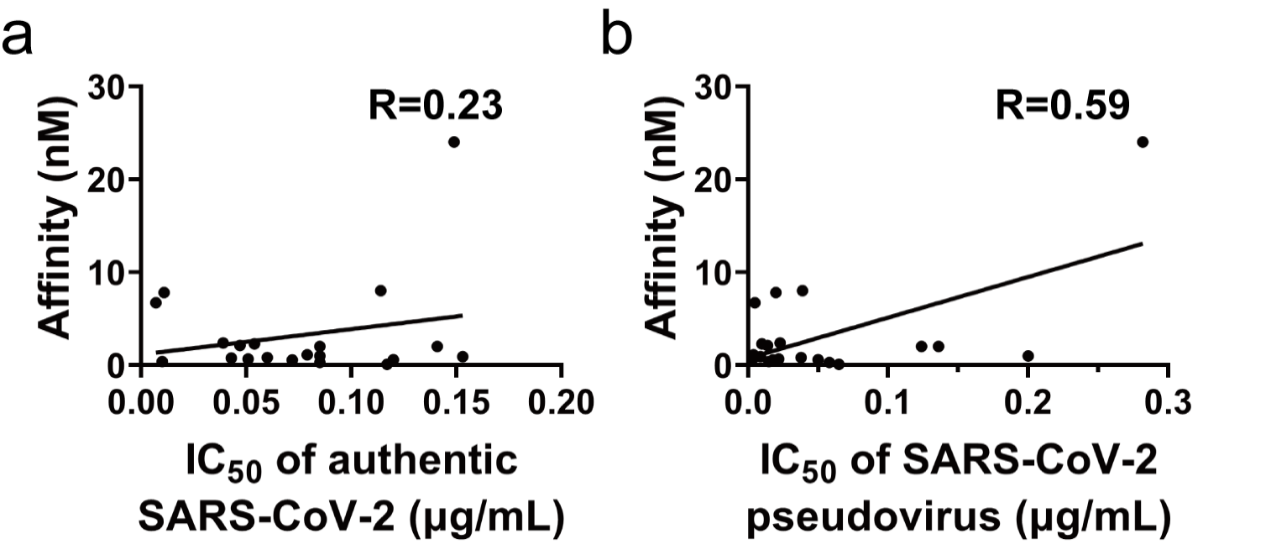

**Extended Data Fig. 6 The correlations of the binding affinities and the IC_50_ of NAbs to block SARS-CoV-2 pseudovirus or the authentic virus.** The correlations between binding affinities and the IC_50_ of NAbs against SARS-CoV-2 pseudovirus (a) or authentic SARS-CoV-2 (b) were analyzed by Pearson Correlation. R^2^ values from the linear regression analysis are shown.

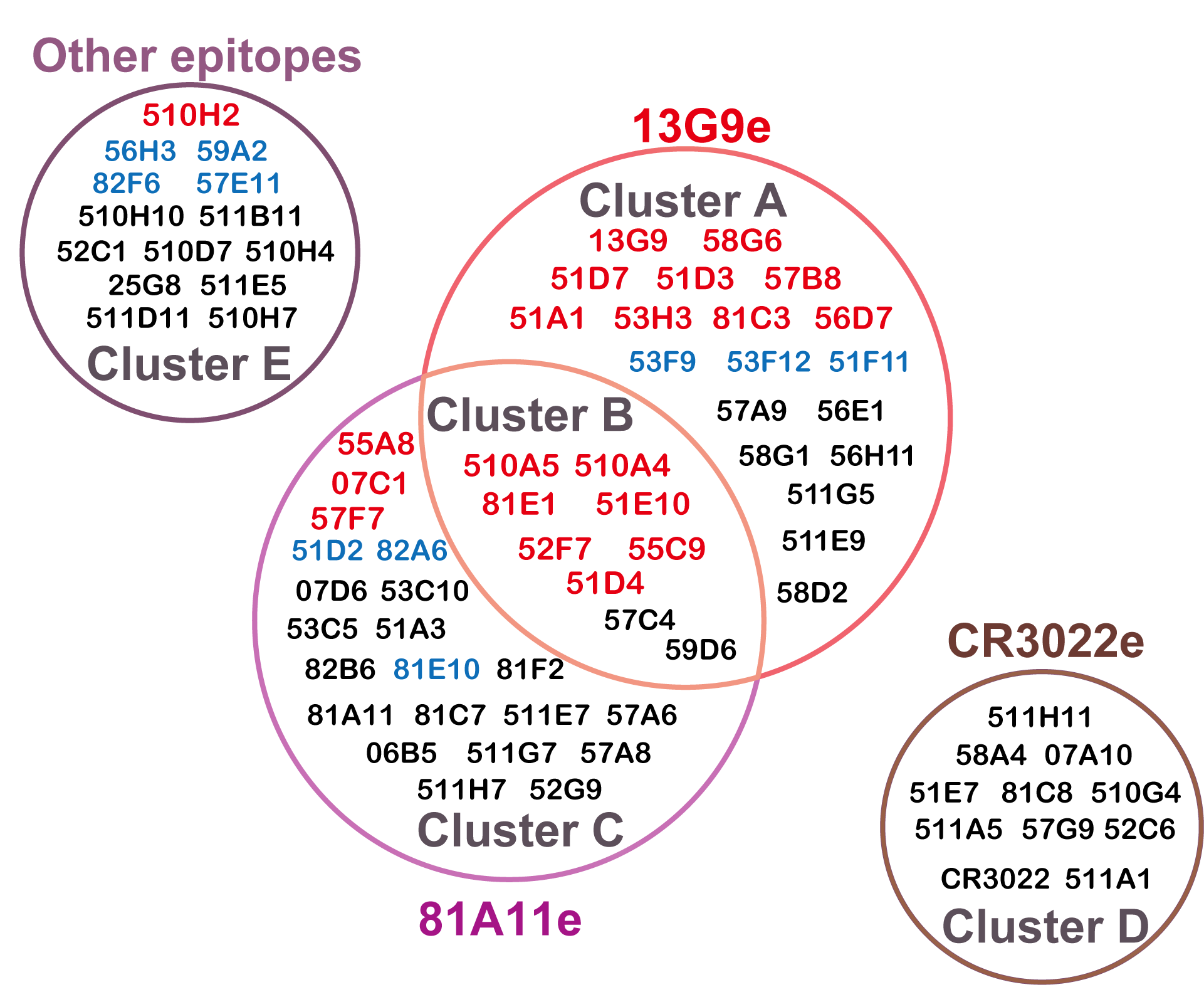

**Extended Data Fig. 7 The interrelationships of mAb clusters were shown in venn diagram.** Different mAb clusters were distinguished by color. The top 20 NAbs were labeled in red and all other NAbs were indicated in blue. The epitopes recognized by the mAbs from corresponding clusters were marked by each circle.

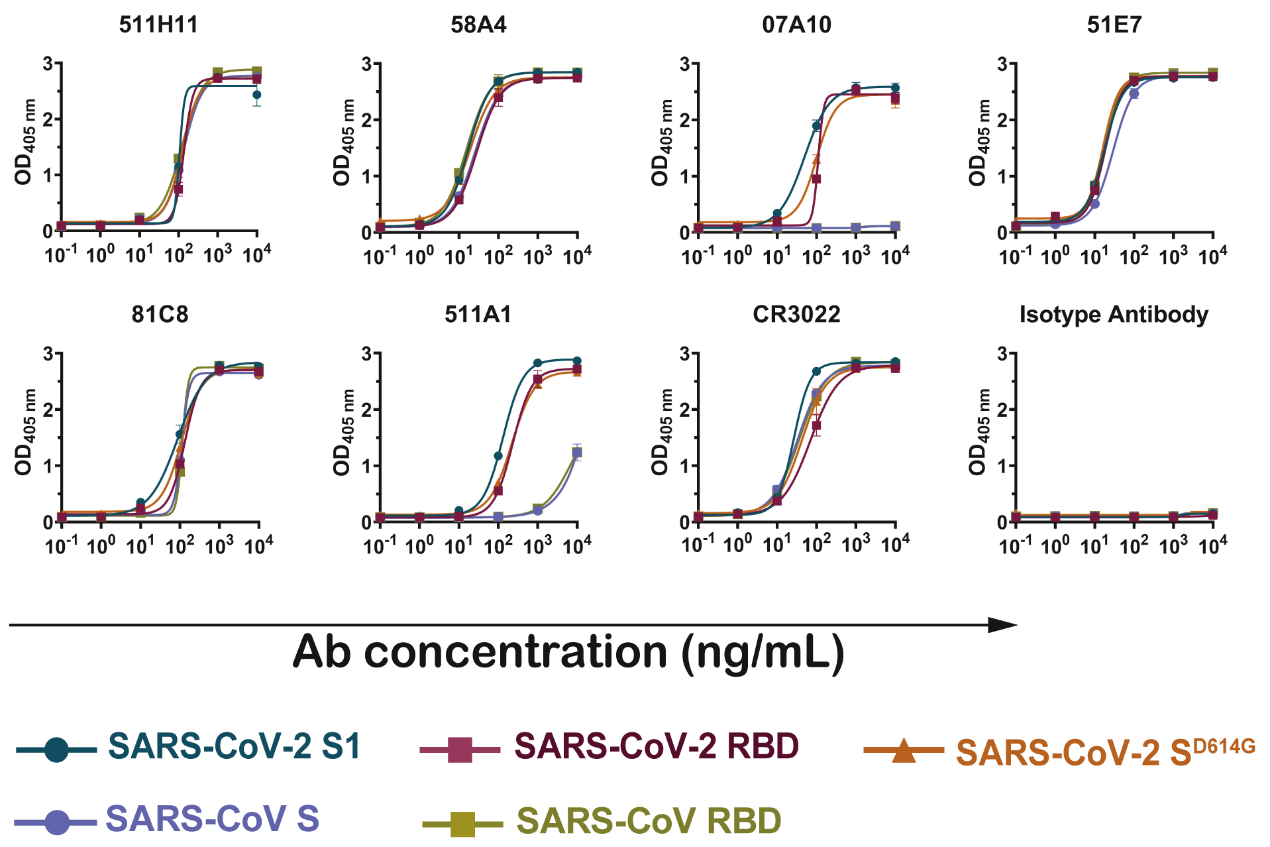

**Extended Data Fig. 8 The binding capabilities of mAbs to SARS-CoV S and S-RBD.** The mAbs in Cluster A could bind to SARS-CoV S and S-RBD, in a concentration dependent manner. CR3022 and the HBV antibody were tested as the positive control and the negative control, respectively. Data are representatives of at least 2 independent experiments performed in technical duplicate. The mean ± SEM of duplicates are shown.

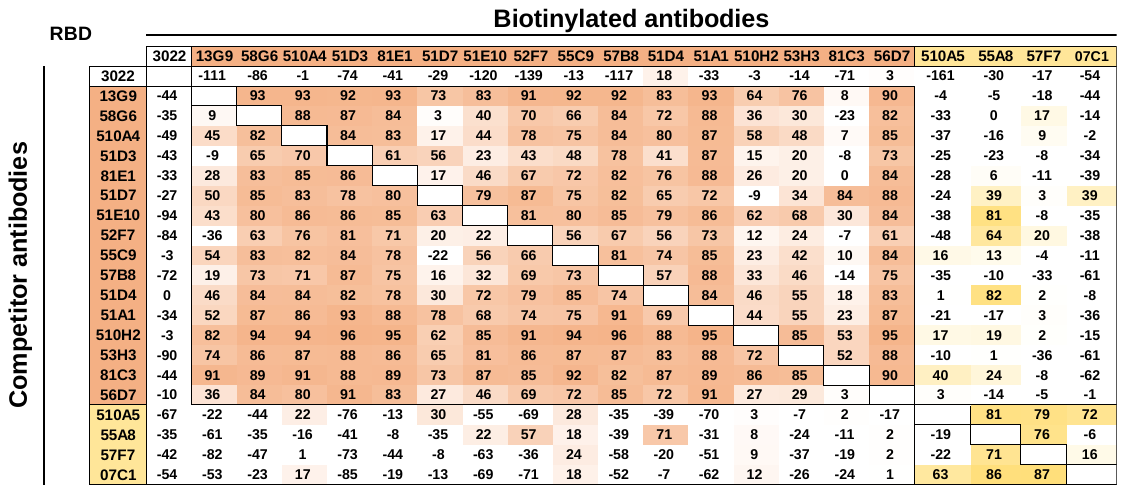

**Extended Data Fig. 9 Competitive study with the top 20 NAbs.** Each NAb was biotinylated, and competed with all other NAbs in the top 20 list bearing no modification, to identify precise epitopes. The numbers in each box are the inhibition rate (%) between two competing NAbs, tested by competitive ELISA. Results are representatives of two independent competitive ELISA experiments.

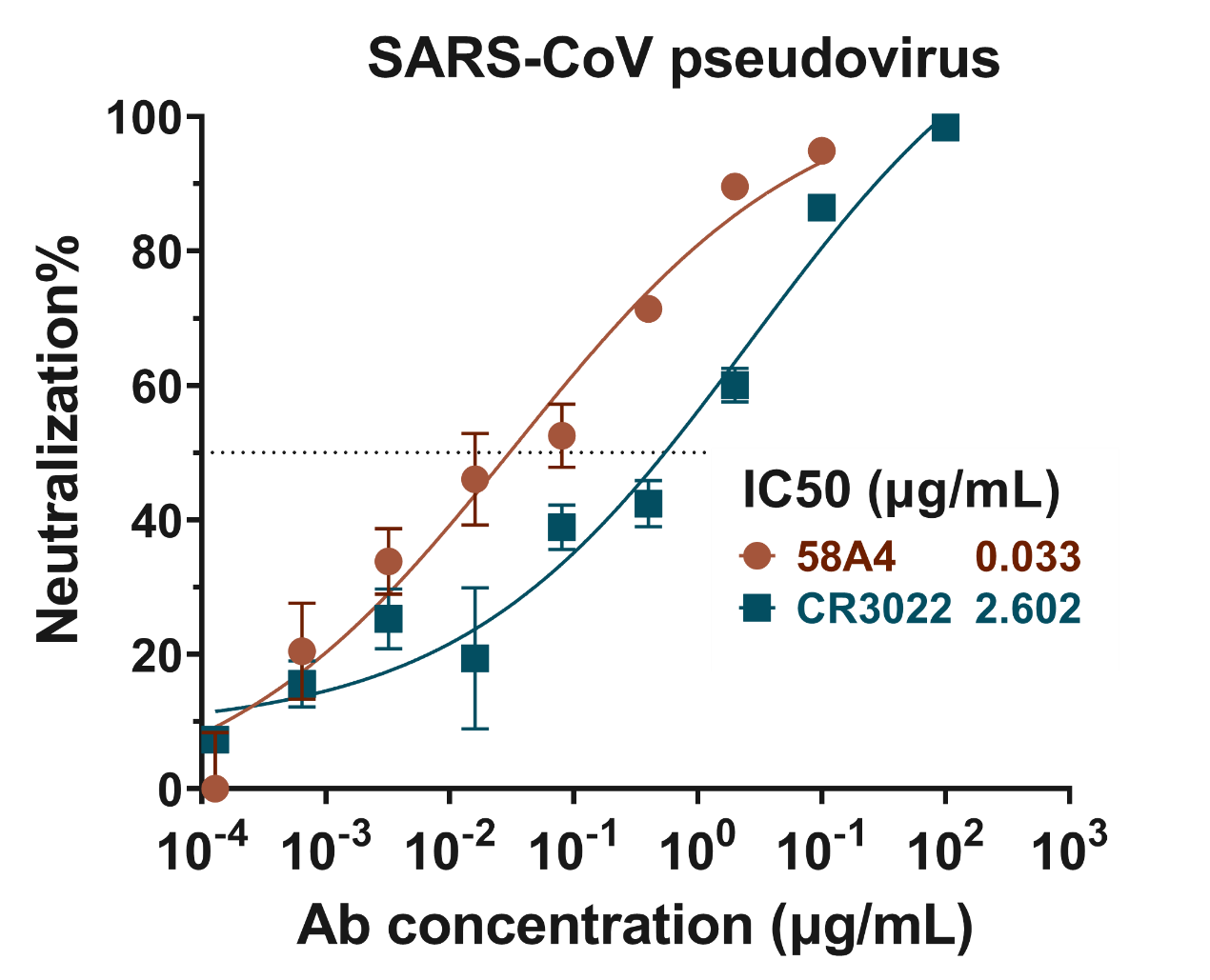

**Extended Data Fig. 10 The neutralization of 58A4 against pseudovirus bearing SARS-CoV.** Serially diluted 58A4 was mixed with SARS-CoV pseudovirus and the neutralizing effects were tested by a luminescence system. The mAb CR3022 that could cross-react with SARS-CoV S-RBD was used as positive control. Data are representatives of at least 2 independent experiments performed in technical triplicate. The mean ± SEM of triplicate are shown.

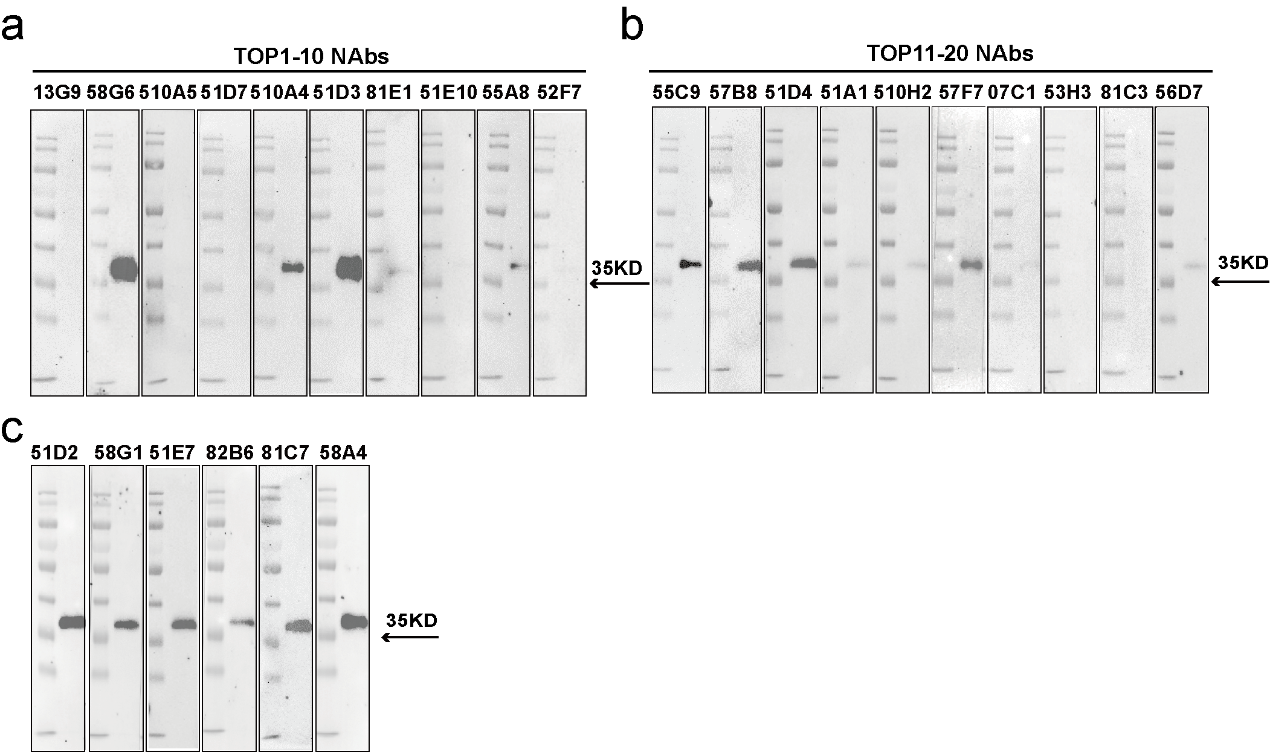

**Extended Data Fig. 11 The binding activities of purified mAbs to the linear S-RBD.** Western blot results of the top 1-10 NAbs (a), 11-20 NAbs (b) and all other mAbs with positive reactions (c) binding to the denatured linear S-RBD. Data are representatives of at least 2 independent experiments performed.

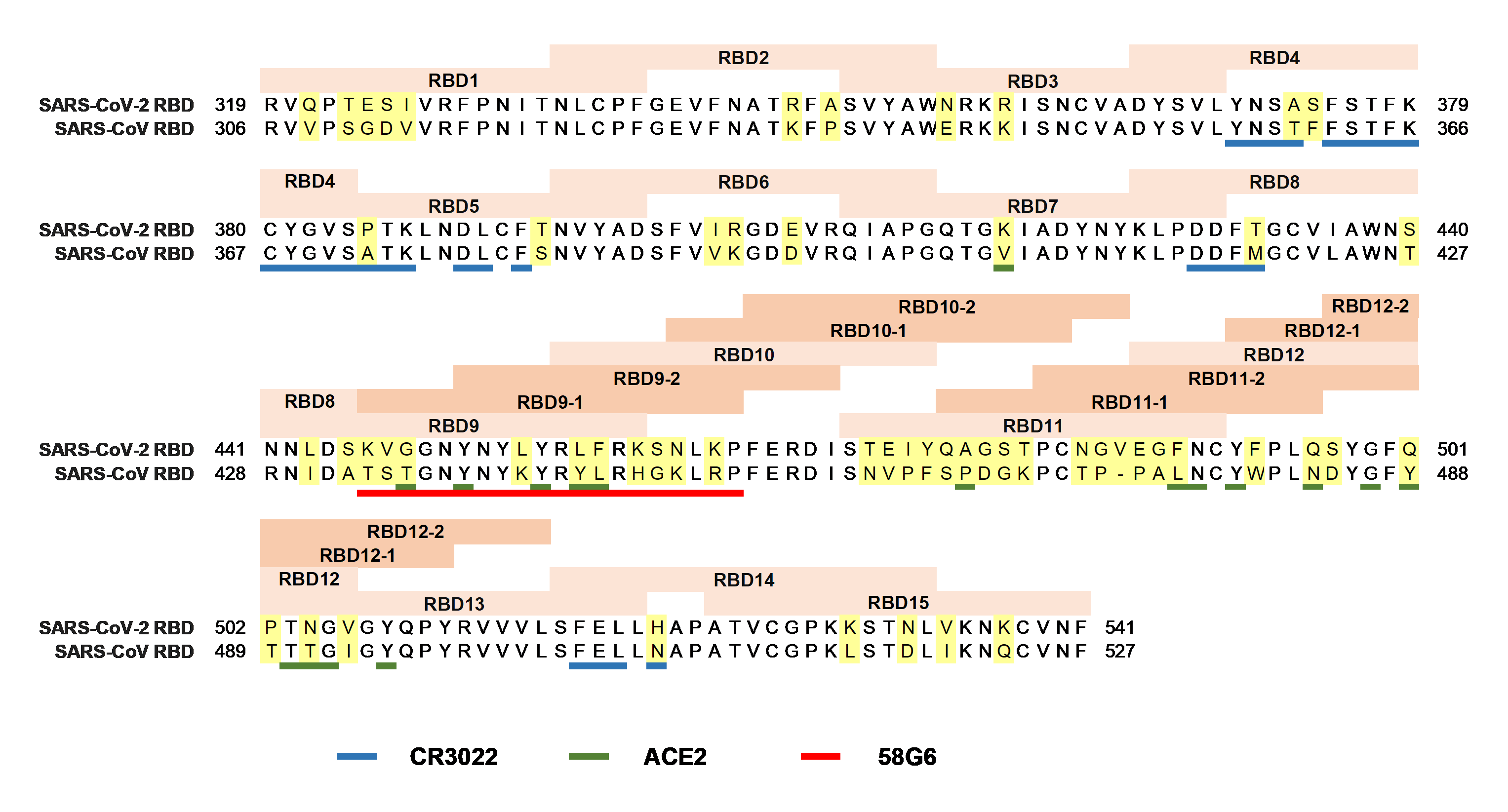

**Extended Data Fig. 12 The profile of peptide design and the antibody binding sites on SARS-CoV-2 S-RBD.** The amino acid sequences of SARS-CoV-2 and SARS-CoV S-RBD were aligned for comparison. Linear peptides designed for SARS-CoV-2 were shown in beige, and the identification of each peptide was indicated in the box above. The non-conserved amino acid residues between two viruses were highlighted in yellow. The binding sites for CR3022, 58G6 and ACE2 were distinguished in different colors as indicated at the bottom.

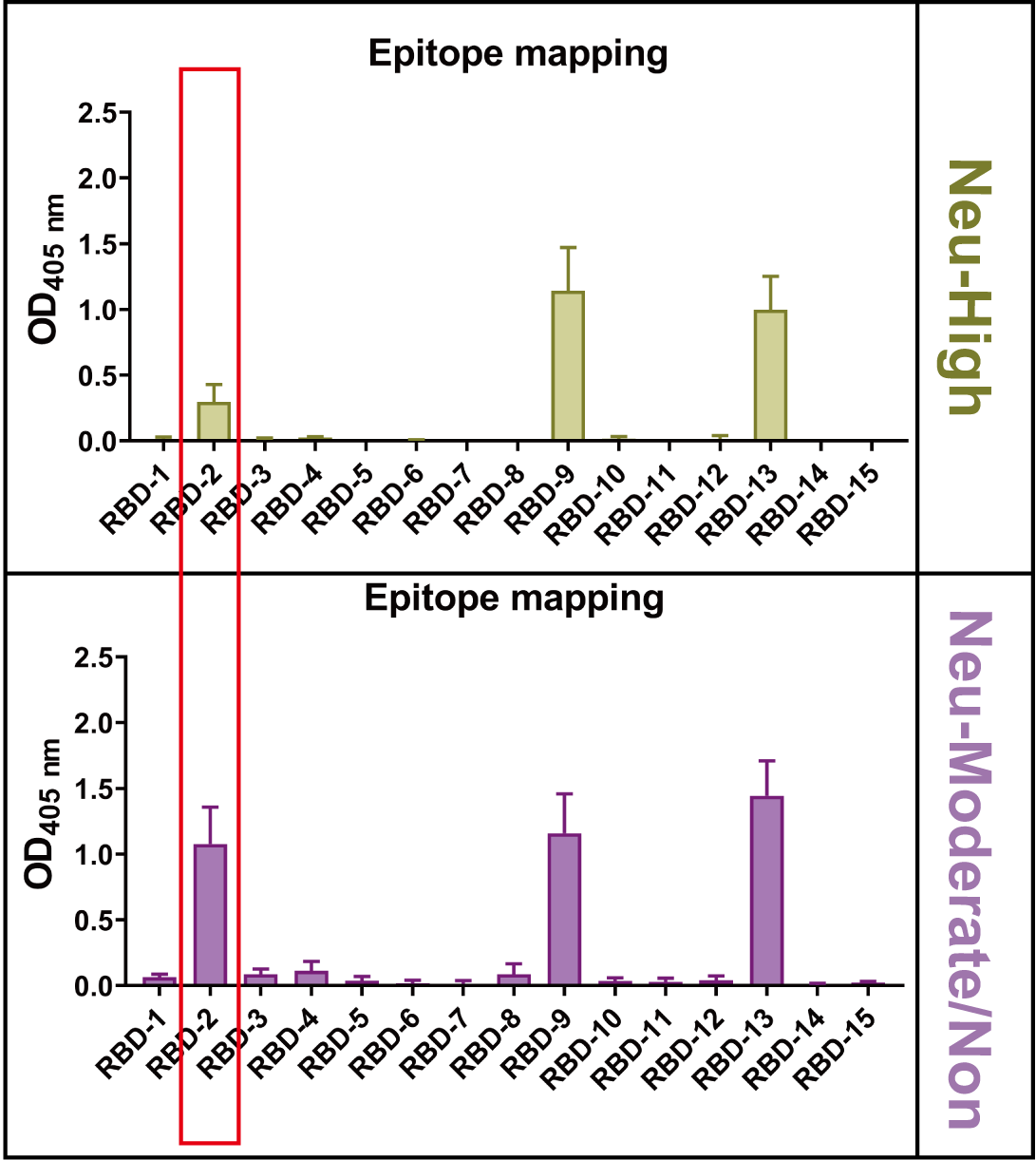

**Extended Data Fig. 13 The mapping of mAbs with linear epitopes represented by designed peptides.** The tested mAbs were divided into two groups according to the neutralizing activities against authentic SARS-CoV-2. 58G6, 510A4 and 51D3 were assigned as Neu-high and all other mAbs were grouped as Neu-Moderate/Non. The interactions of these mAbs to the linear epitopes represented by designed peptides were analyzed by peptide ELISA. Six mAbs did not react with synthesized peptides and not appeared in these groups. The negative correlation of the RBD2 reactivity of these mAbs with their neutralizing capability was marked with a red box. The mean ± SEM of duplicate are shown.

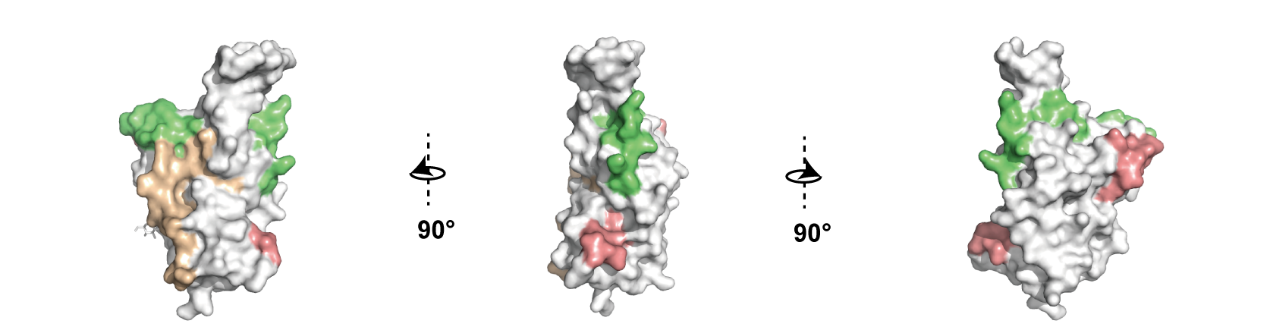

**Extended Data Fig. 14 The positional relations of linear epitopes represented by RBD2, RBD9-1 and RBD13 on SARS-CoV-2 S-RBD.** The structure of SARS-CoV-2-RBD was shown at three depositions and the epitopes represented by RBD2 (yellow), RBD9-1 (green) and RBD13 (pink) were labeled for easy observations of their positional relations.

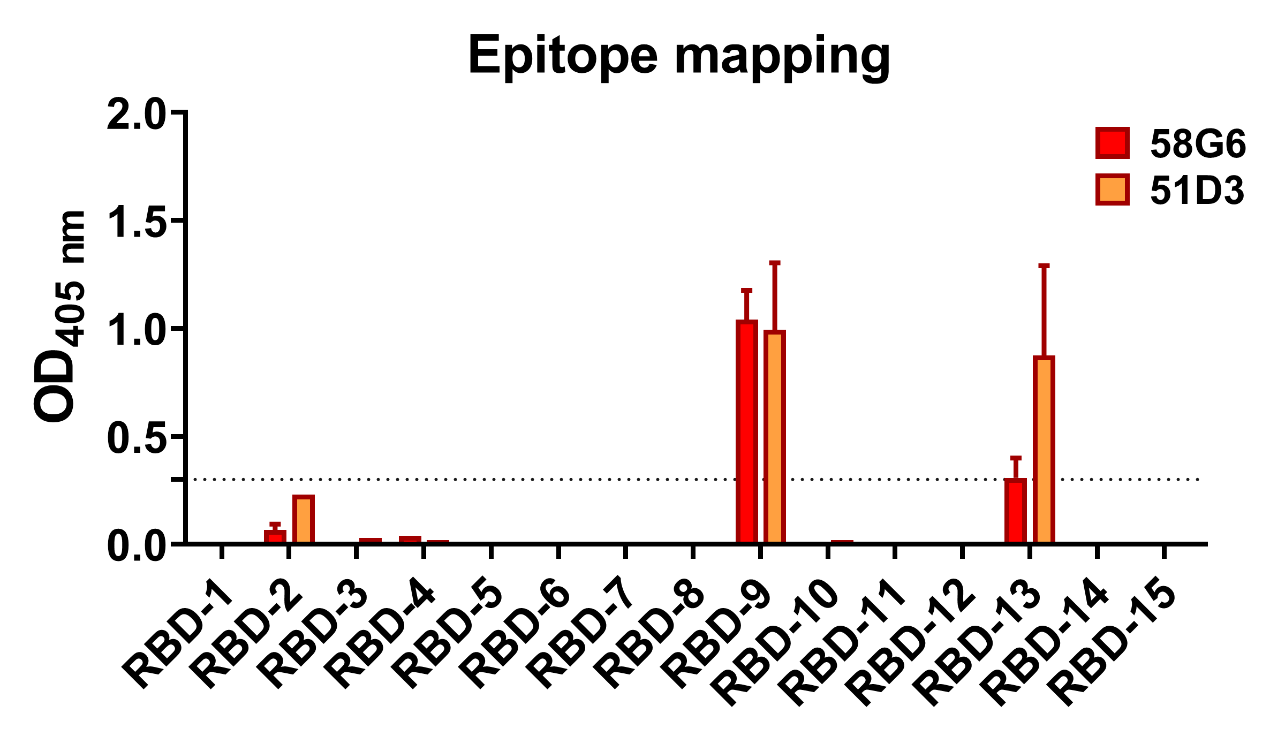

**Extended Data Fig. 15 The binding ability of 58G6 and 51D3 to the linear peptides.** The interactions of 58G6 or 51D3 with the linear peptides were analyzed by peptide ELISA. The dotted line indicates the cut-off level of positive responses (OD>0.3). Data are representative of at least 2 independent experiments performed in technical duplicates. The mean ± SEM of duplicates are shown.

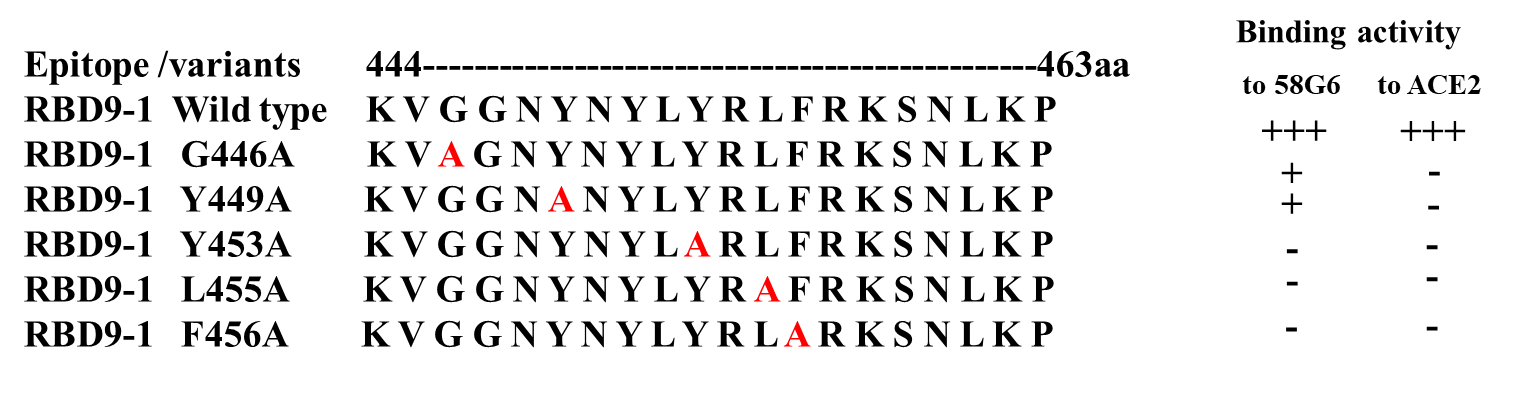

**Extended Data Fig. 16 The sequences of RBD9-1 and its point mutations.** Summary of the binding activities of mutated RBD9-1 peptides to 58G6 that were shown in fig 4a, or those to ACE2 that were shown in 4b. The mutated amino acids of RBD9-1 were indicated in red.

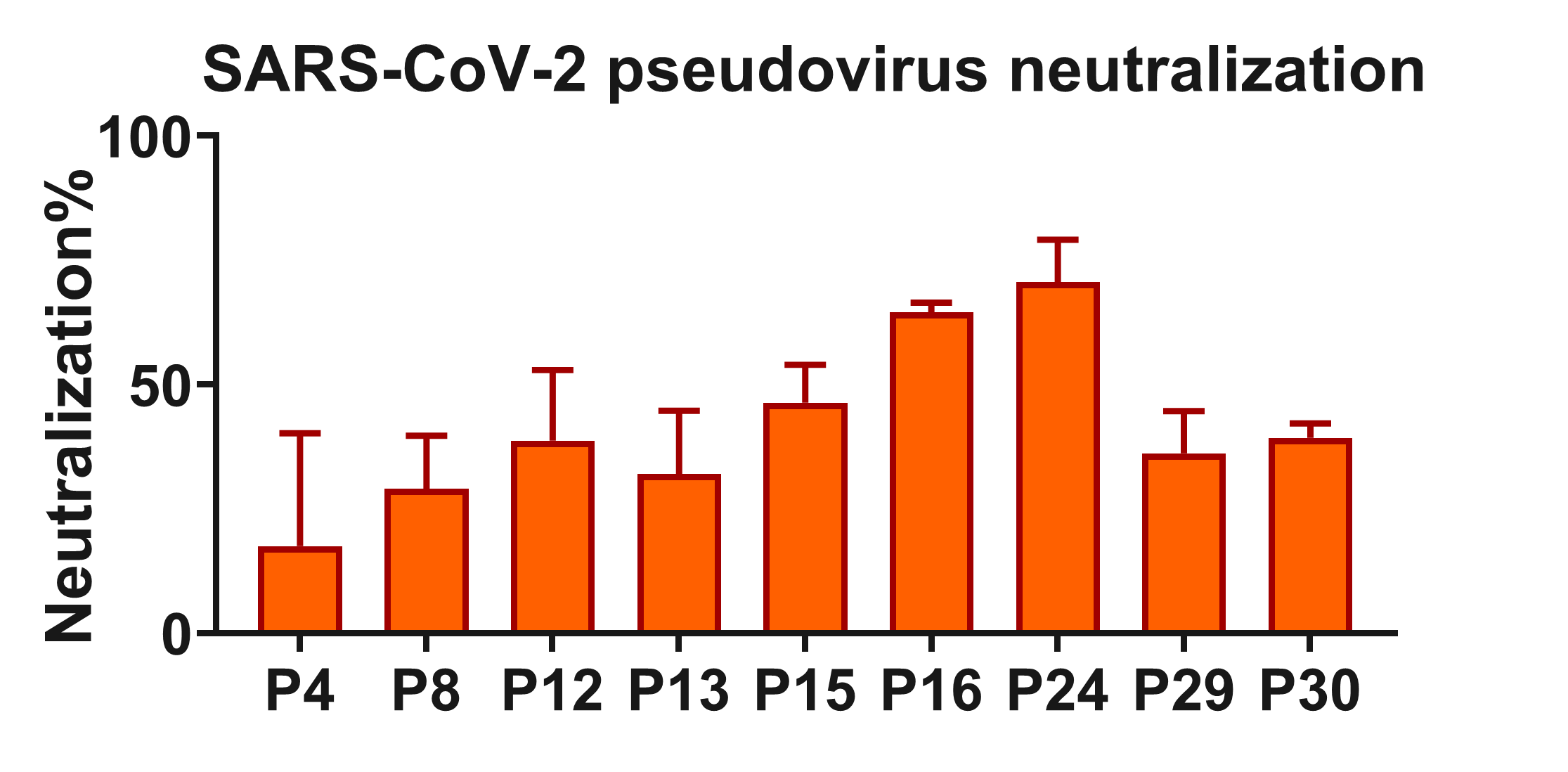

**Extended Data Fig. 17 The neutralizing capabilities of convalescent patients’ plasma against SARS-CoV-2 pseudovirus.** The COVID-19 convalescent patients’ plasma (1:1000) was mixed with SARS-CoV-2 pseudovirus, and the neutralizing effects were tested by the luminescence system. Data for each NAb were obtained from a single experiment, with three technical replicates. Data are presented as mean ± SEM.

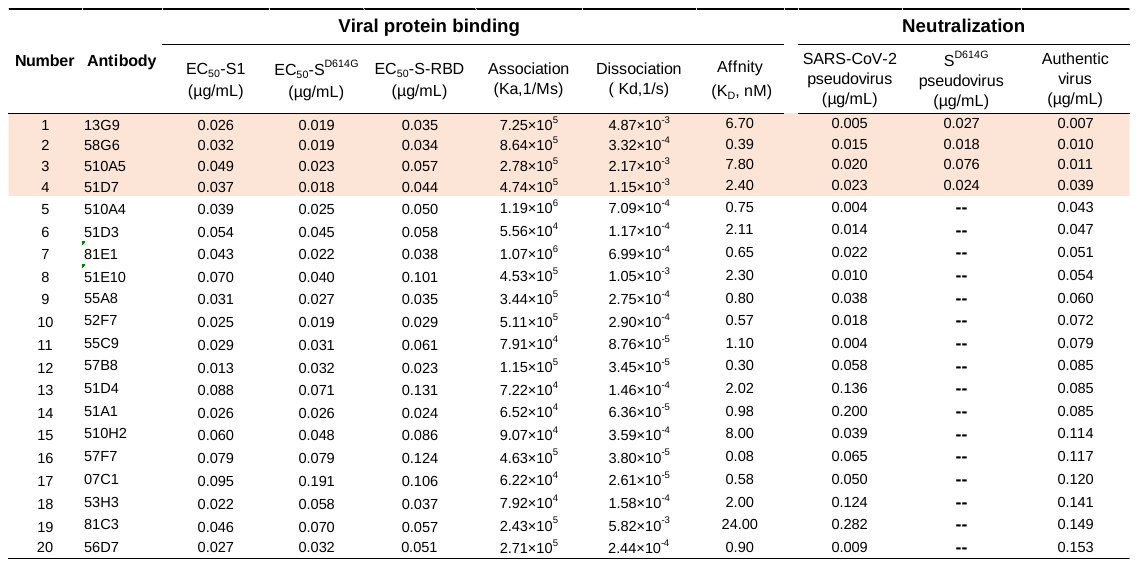
Table 1. The characteristics of the top 20 NAbs

The top 4 NAbs with the most potent neutralizing capabilities were highlighted in orange. --: not analyzed.
